# Species-conserved mechanisms of abstract rule learning promote cognitive flexibility in complex environments

**DOI:** 10.1101/2022.11.14.516439

**Authors:** Florian Bähner, Tzvetan Popov, Nico Boehme, Selina Hermann, Tom Merten, Hélène Zingone, Georgia Koppe, Andreas Meyer-Lindenberg, Hazem Toutounji, Daniel Durstewitz

## Abstract

Rapid learning in complex and changing environments is a hallmark of intelligent behavior. Humans achieve this in part through abstract concepts applicable to multiple, related situations. It is unclear, however, whether some of the underlying computational mechanisms also exist in other species. We combined behavioral, computational and electrophysiological analyses of a multidimensional rule-learning paradigm in rats and humans. We report that both species infer task rules by sequentially testing different hypotheses, rather than learning the correct action for all possible combinations of task-related cues. Neural substrates of hypothetical rules were detected in prefrontal network activity of both species. This species-conserved mechanism reduces task dimensionality and explains key experimental observations: sudden behavioral transitions and facilitated learning after prior experience. Our findings open the black box of hypothesis testing in rodents and provide a foundation for the translational investigation of impaired cognitive flexibility.

## Introduction

Cognitive flexibility is critical for the ability to respond to changes in the environment in adaptive ways. Deficits in this domain are observed in several major neuropsychiatric disorders.^1^ However, it is not fully understood how correct behavioral rules are identified in ever-changing contexts and how this information is encoded in neural activity.^2–7^ A powerful theoretical framework in this area is reinforcement learning (RL) which describes how action values in different environmental situations (known as states) are learned to guide decisions and maximize reward.^8^ This framework has gained support from the neurosciences following the discovery of neural substrates of RL quantities such as action values (i.e., the expected reward from taking that action) and reward prediction errors (i.e., the difference between actual and expected reward).^9,10^ A core assumption of classic RL models is that subjects learn to map each state of the environment to the maximum-value action.^8^ However, the real world is multidimensional, where many sources of information, such as sensory cues, reward history, and working memory of past choices combine to form each state. This results in an exponential growth in the number of states and state-action mappings (known as “curse of dimensionality”) that translates into slow and gradual learning.^11^ Consequently, standard RL models struggle to explain hallmarks of flexible behavior such as sudden transitions in performance^7,12–15^, rapid learning in complex environments, or faster learning with prior experience.^3,11,16^

A long tradition in cognitive science maintains that humans work around the curse of dimensionality by learning abstract concepts like rules,^3,17^ categories^18,19^ or schemas^20,21^ that can be applied to multiple, related situations. Recent advances in artificial intelligence inspired by cognitive neuroscience have suggested different computational mechanisms for learning abstract models of the world that facilitate generalization of knowledge and transfer of learned skills to new tasks.^3,16,22–26^

Since one-to-one mappings of states to actions are inefficient in high dimensional environments with many potential task rules, subjects may rather learn and test *hypothetical rules* underlying these mappings, such as *“don’t press any lever paired with a bright light, regardless of lever location”*. This abstraction reduces dimensionality dramatically since every cue becomes an instance of a general task feature, allowing the learner to direct its attention toward this feature and ignore other cues. The learner can then test different hypothetical rules against environmental evidence until they successfully identify the experimenter-defined rule. We call such hypothetical rules *behavioral strategies* in the following to distinguish them from task rules.

Indeed, flexible human behavior can be explained by computational models that are compatible with the above framework, including hierarchical and attention-modulated RL models.^3,5,16,27^ It is unknown, however, how rodents infer task rules in complex environments and whether some of the underlying behavioral and neural mechanisms are conserved across species. Pioneering work has shown that in some contexts, rats do follow behavioral strategies like *win-stay-lose-shift* or *spontaneous alternation*^28,29^ and recent methodological work has shown that such strategies can be identified in rodents, macaques and humans.^30^ However, these findings have not been generalized to cognitive flexibility or empirically contrasted to the prevailing view of classical RL models.

We developed a novel translational rule-learning task and used a combination of behavioral, computational, and electrophysiological methods to test whether common computational mechanisms exist in both species. We found that both rats and humans followed low-dimensional behavioral strategies to test each task feature for relevance rather than learning all possible mappings between environmental states and actions simultaneously. Moreover, we decoded strategy-related quantities with features of abstract, hypothetical rules from both rat multiple-single unit and human magnetoencephalography (MEG) recordings in the prefrontal cortex (PFC), highlighting a common mechanism of flexible rule learning.

## Results

### Rats infer task rules using strategies

We developed a novel dual-choice multidimensional rule-learning paradigm with four relevant task features to evaluate the hypothesis that rats identify experimenter-defined rules by sequentially testing different low-dimensional behavioral strategies (Figure 1A). A loudspeaker and a cue light were positioned above each lever. In each trial, one auditory and one visual cue were presented to model multisensory input. Rats were not pre-trained on any rule and had to learn one out of eight possible rules with deterministic reward feedback: go-click (i.e., follow the auditory cue in the presence of a distracting visual cue), go-silent, go-light, go-dark, go-right, go-left, alternate or win-stay-lose-shift/WSLS. Thus, four task features could be relevant for reward in every trial: the visual and auditory cues as well as outcome and choice histories. Rats were able to learn each rule, albeit at different learning rates (Table S1). We used change point analysis and behavioral modeling to identify when performance increases occurred during rule learning and how abrupt they were (Methods).^31,32^ We detected transitions that are best characterized as a step change (median (Q_1_-Q_3_) of 10-90% rise time: 1 (1-1) trials, n=117) similar to what has been observed in other, simpler learning paradigms.^7,12,13^ This finding is at odds with the classic RL view that assumes learning to be slow and gradual since rats would need to learn the correct action (e.g., right or left lever press) for each of the 2^4^=16 states (see Figure 1B, top for one example). We therefore propose an alternative hypothesis (Figure 1B, bottom) where rats use up to eight low-dimensional behavioral strategies (*go-light/go-dark*, *go-click/go-silent, go-right/go-left*, *alternate, or win-stay-lose-shift*) to sample different features for relevance (i.e., significance of sensory cues, past choices and outcome history). In this scenario, rats would test different simple hypothetical rules sequentially until they infer the correct task rule.

**Figure 1.**
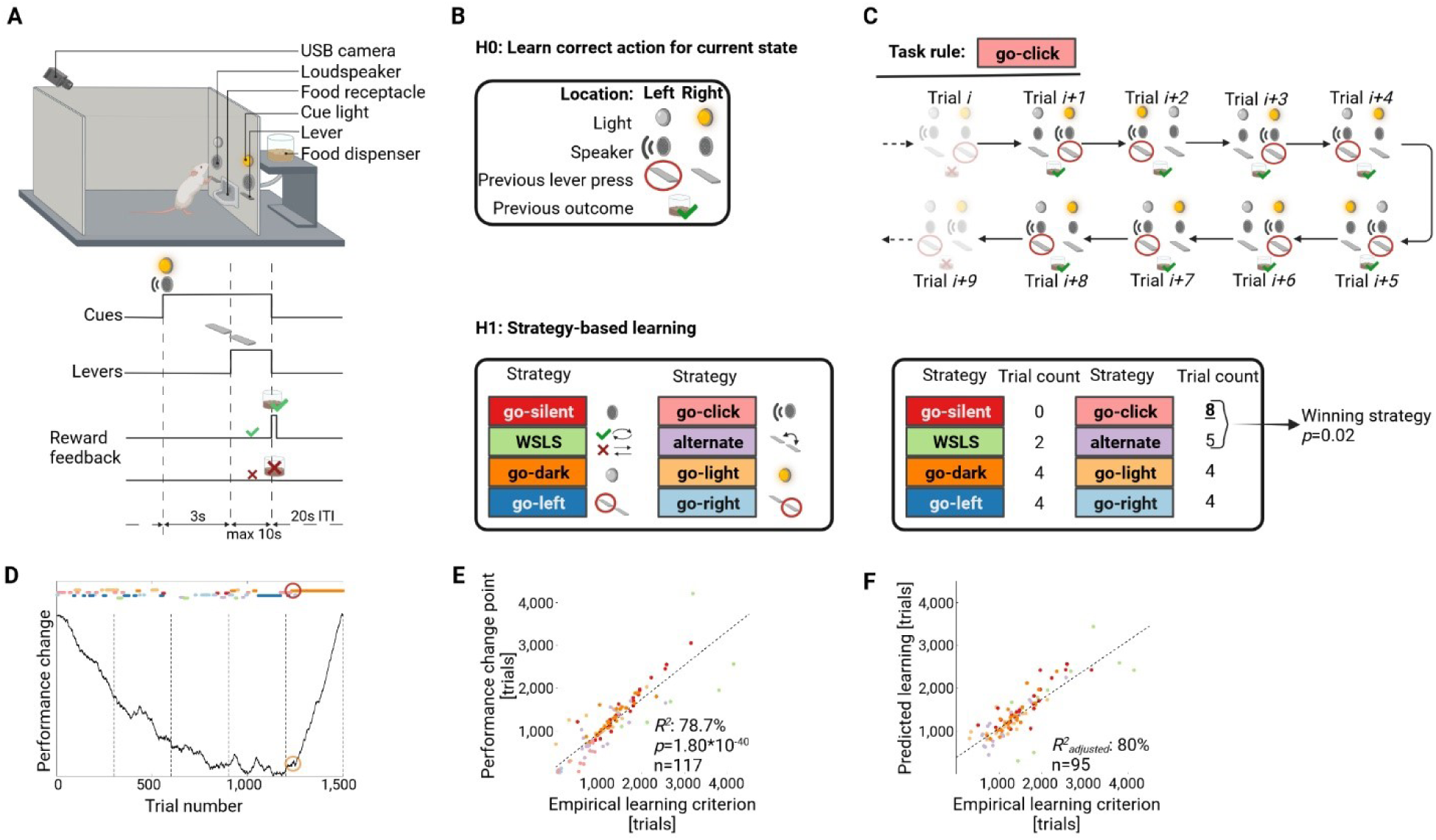
Rats test behavioral strategies to infer the task rule. (A) Each trial starts with the presentation of one auditory and one visual cue. Three seconds later, two levers extend and rats use trial-and-error to learn the correct choice. Depending on the experimental rule, one out of four task features can be relevant for reward. (B) Our core hypothesis is that rats sequentially follow low-dimensional behavioral strategies (*bottom*) to test these four task features for relevance (e.g., *go-click*/*go-silent* whether the sound is relevant). The null hypothesis assumes that animals learn all possible mappings between cue combinations (see *top* for example) and actions. (C) *Top:* example of a *go-click* sequence consisting of eight consecutive trials as detected with the strategy detection algorithm. For each trial, the current sensory input is shown as well as the choice and reward feedback for this trial. *Bottom:* the count of trials that are consistent with each strategy for the highlighted block of eight trials is shown. The binomial statistic based on this count indicates that that the rat followed the strategy *go-click*. (D) Performance change of an individual rat learning the task rule go-dark. Strategy sequences are color-coded, performance change is visualized using a CUSUM (cumulative sum of differences to the mean) plot. A global minimum in the CUSUM curve indicates that performance increases with respect to mean performance after the change point. Detected performance change point and empirical learning criterion are marked with circles, dashed vertical lines mark the end of an experimental session. (E) High correlation between empirical learning criterion and performance change point, each dot is a rat learning one out of seven different rules. (F) A linear regression model shows that strategy-related parameters significantly predicted the learning trial. See Figure S1, Table S2. ITI/inter-trial interval.

To test this scenario, we developed a strategy detection algorithm that measures whether a particular strategy is more likely than chance or any other strategy within every possible trial window (Methods, Figure 1C). The algorithm identifies trial sequences within each session in which a particular strategy is the winning strategy. In each task rule, animals indeed followed a median of 7 (6-8) out of 8 possible strategies (Table S1). The average length of an individual strategy sequence (i.e., the number of consecutive trials where a strategy was followed) was 14 (12-16.3) trials. Interestingly, we replicated this finding in a separate cohort of rats (n=19) that were presented with random reward feedback (i.e., none of the four task features were predictive of reward), indicating that hypothesis testing reflects a general cognitive strategy, even when there is no behavioral advantage. We defined an empirical measure for learning based on the comparison between the length of a correct strategy sequence (i.e., the strategy that corresponds to the current task rule) and the distribution of sequence lengths of that same strategy as detected in the cohort that received random reward. For example, we assume a rat learned the rule go-silent successfully when the sequence length of the corresponding *go-silent* strategy was at least 21 trials. Beyond this threshold, *go-silent* sequences were considered outliers in the random reward cohort (Methods, Table S1). The trial at which rats reached the empirical learning criterion correlated significantly with abrupt performance increases (Figures 1D, 1E, and S1). Furthermore, we reasoned that if rats follow strategies to identify task rules, more efficient strategy use should lead to faster learning. Indeed, multiple regression analysis revealed that several strategy-associated quantities were predictive of the trial at which the empirical learning criterion was reached (R^2^_adjusted_=80%, Figure 1F), including the trial number at which the animal followed the correct strategy for the first time (p=1.6*10^-4^), the number of correct strategy sequences before reaching the empirical learning criterion (p=5.0*10^-10^), a perseveration measure^33^ at the strategy level (p=9.0*10^-20^), and r-1 binary indicator variables coding r task rules (Figure S1, Table S2). While not all trials (36.16 ± 1.24%) could be assigned to a specific strategy, the number of such trials was not predictive of learning since adding it to the regression model did not increase goodness of fit (R^2^_adjusted_=78.9%, p=0.43 for this regressor). We found comparable results in rats (n=16) performing a classic operant two-rule set-shifting task^33^ where rats also followed behavioral strategies that predicted sudden transitions in performance (Figure S2). In sum, this analysis provides first indication that to infer task rules, rats test low-dimensional behavioral strategies rather than learning high-dimensional mappings between environmental states and actions.

### Strategy-specific selective attention mechanisms in rats

Selective attention focused on different task features is a candidate mechanism for guiding behavioral responses (i.e., the rat focuses on one general task feature at a time instead of a combination of low-level cues). This could explain why rats follow strategies for many trials, even when reward is random. Selective attention is also able to reduce dimensionality in complex environments to a manageable subset of task features that can be tested for relevance.^5,11^ To test the role of attention in our paradigm, we applied machine learning techniques (Methods) to screen videotaped sessions for behavioral markers of attention in another group of rats that performed multiple consecutive rule switches (n=29 rats). We found a strong orientation reaction at cue onset that was influenced by the current strategy. More specifically, we found that the onset and offset trials of each strategy sequence were locked to abrupt changes in head direction at cue onset towards and away from the task feature that is relevant for that strategy (Figure 2A). This allowed us to obtain a trial-by-trial measure of the rat’s attention to each task feature at the time of cue presentation, which we term “attention-at-choice”. Strategy-specific abrupt changes in attention to a corresponding task feature were found for all eight strategies (Figure S3). We observed a similar, but feature-independent, orientation reaction during reward feedback, which we term “attention-at-reward” (Methods). These findings were replicated in the random reward cohort (n=19, Figure S4). Rats thus test different hypotheses sequentially, even when they have no incentive to follow a specific strategy.

**Figure 2.**
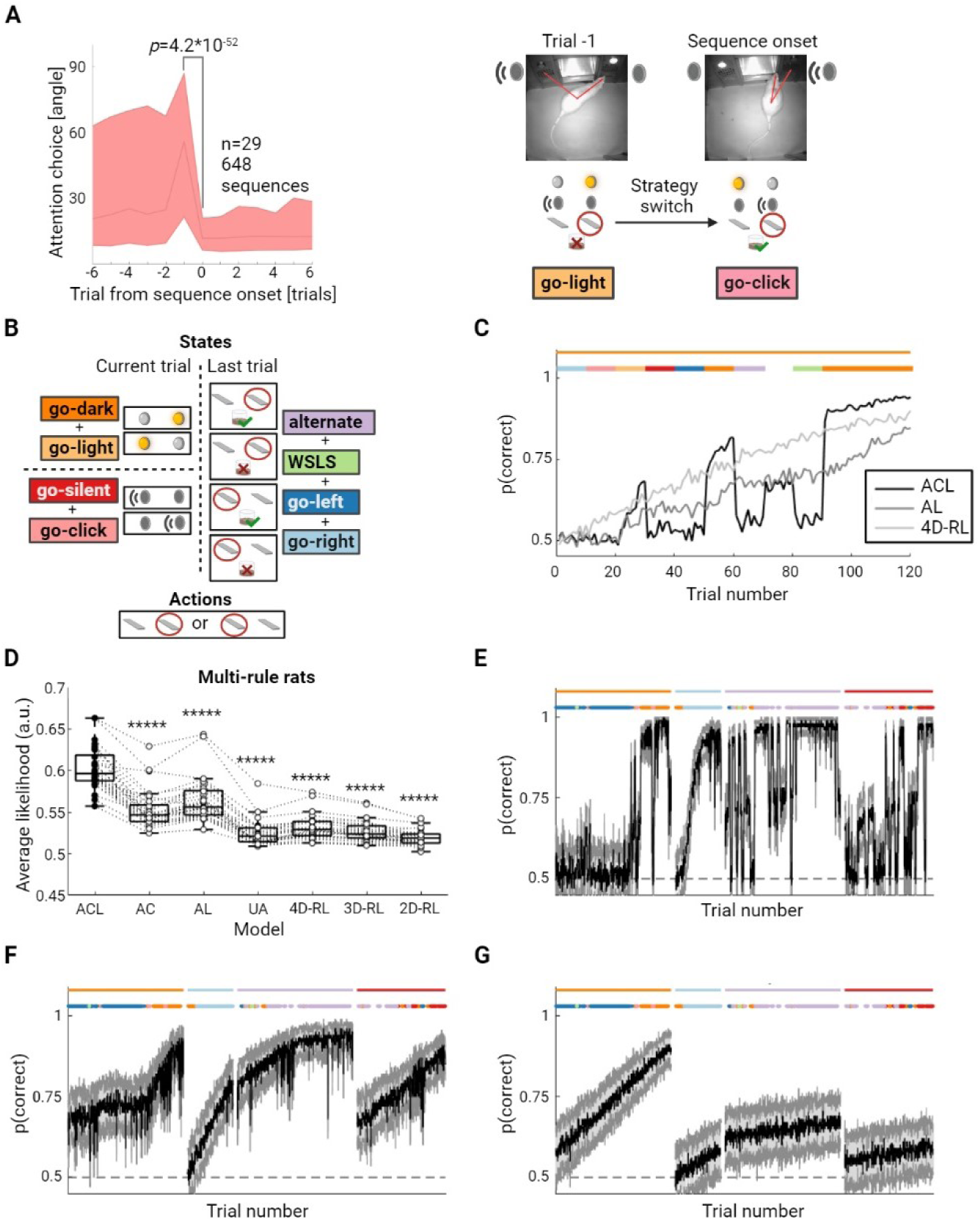
Strategy-specific attention during choice and learning. (A) *Left:* Head direction plot for strategy *go-click* during choice formation. Head angles with respect to sound are aligned to cue onset and abruptly decrease at sequence onset (trial 0). See Figures S3, S4. *Right:* example of head angles in trial before and at sequence onset. (B) Visualization of the state-action space used in attention-based RL models. Strategies map to distinct state spaces via their attention focus. (C) Example learning curves of RL agents (gray-scale lines) trained on the task go-dark (solid orange line), averaged over 1000 simulations. The agents switch their attention between different strategies, before settling on the correct strategy (filled colored circles). Attention score of the attended to state subspace is set to 0.8. All models used the same parameters: learning rate alpha=0.2; exploration rate beta=3. Step changes are only observed for models that have an attention-at-choice component. (D) Trial-averaged cross-validated likelihood for seven RL model variants. Whiskers: 1.5 IQR; dashed lines connect values (circles) of the same rat; filled circles represent best (i.e., highest likelihood) model of each rat. The ACL (attention biases both choice and learning) model is the winning model in each rat. Model comparisons between ACL and all other models are Benjamini-Hochberg corrected (Wilcoxon matched pairs test on non-normalized BICs with *****: p<10^-5^). (E-G) Representative example of model simulations using model parameters estimated from the experimental data of one rat. Step changes in performance could only be reproduced if choice is modulated by the strategy-specific attention focus (ACL in E), but not AL (F) or 4D-RL models (G). Task rules and strategies are color-coded as in Figure 1.

To assess whether the two sets of attention measures are relevant for multidimensional rule learning, we incorporated them into different RL models (Methods) to evaluate how rats learn rules in the state-action space defined by the four task features (where the state space consists of three subspaces: a visual, an auditory, and a 2-dimensional history state combining the place and outcome features; Figure 2B). More specifically, we tested whether attention modulates behavior during choice alone (attention-at-choice/AC model), learning (or reward feedback) alone (attention-at-learning/AL model), or both (ACL model).^11^ The attention measures bias learning and choice in the ACL model such that the task feature related to the current strategy has a privileged position. In a fourth model, attention is split equally between all three subspaces so that it neither biases choice nor learning (uniform-attention/UA model). We also compared these attention-modulated RL models to three variants of a standard RL models that mapped a multidimensional state space directly to actions. These standard models differed in which task features comprised the relevant states: 1) a 4-dimensional RL model (4D-RL) where the state space consisted of all 4 relevant task features, resulting in 32 state-action mappings to be learned, 2) a 3-dimensional RL model (3D-RL) that excluded the outcome feature, and 3) a 2-dimensional RL model (2D-RL) that only included the sensory task features.

We first compared the models through simulations of artificial agents that aimed to learn an experimenter-defined rule (e.g., go-dark). We assumed this artificial agent switched their attention between all eight possible strategies every 10 trials, including a period of 10 trials where it attended to no strategies at all, before settling on attending to the correct task feature (e.g., the light cue) for 30 trials. We found that attention during choice (ACL and AC models) is necessary to produce sudden transitions in the agent’s behavior as it learned the correct task rule. In contrast, models where attention modulated learning alone (AL model) or where the agent learned all or partial mappings between states and actions (UA model and standard RL models) resulted in slow and gradual learning (Figure 2C).

Next, we fitted the parameters of the four attention-modulated RL models and the three standard RL models to experimental data, followed by cross-validated model comparison. This revealed that the ACL model was indeed the best at predicting held-out behavioral data in all 29 rats (Figure 2D). We further confirmed that attention at choice is necessary to capture sudden improvements in performance, a core feature of rat multidimensional rule learning: we ran model simulations using model parameters as estimated from the experimental data and constrained by sensory cues and attention scores as measured experimentally. We found that step changes in performance similar to what is observed for experimental data could only be reproduced if choice is modulated by the strategy-specific attention focus (Figures 2E-G).

In summary, our data suggest strategy-specific selective attention as a mechanism to speed up learning by reducing task dimensionality in complex environments, since it allows animals to focus on one task features at a time, rather than dividing attention between specific cues.

### Formation and value-based selection of behavioral strategies in rats

So far, our analyses show that strategy-specific attention constrains action selection in response to a stimulus but cannot explain how strategies and attention focus are selected in the first place. A different class of RL models used to explain human rule learning^3,27^ suggests that additional computational processes are at work. More specifically, action selection is proposed to be hierarchical such that strategies are hypothetical rules operating as high-level, abstract actions that are selected first (e.g., *go-light*) and then map specific stimuli (e.g., cue light left) to low-level actions (e.g., left lever press; Fig. 3A). Such hierarchical RL models make several behavioral and neural predictions^3,34,35^ we tested empirically using our paradigm.

**Figure 3.**
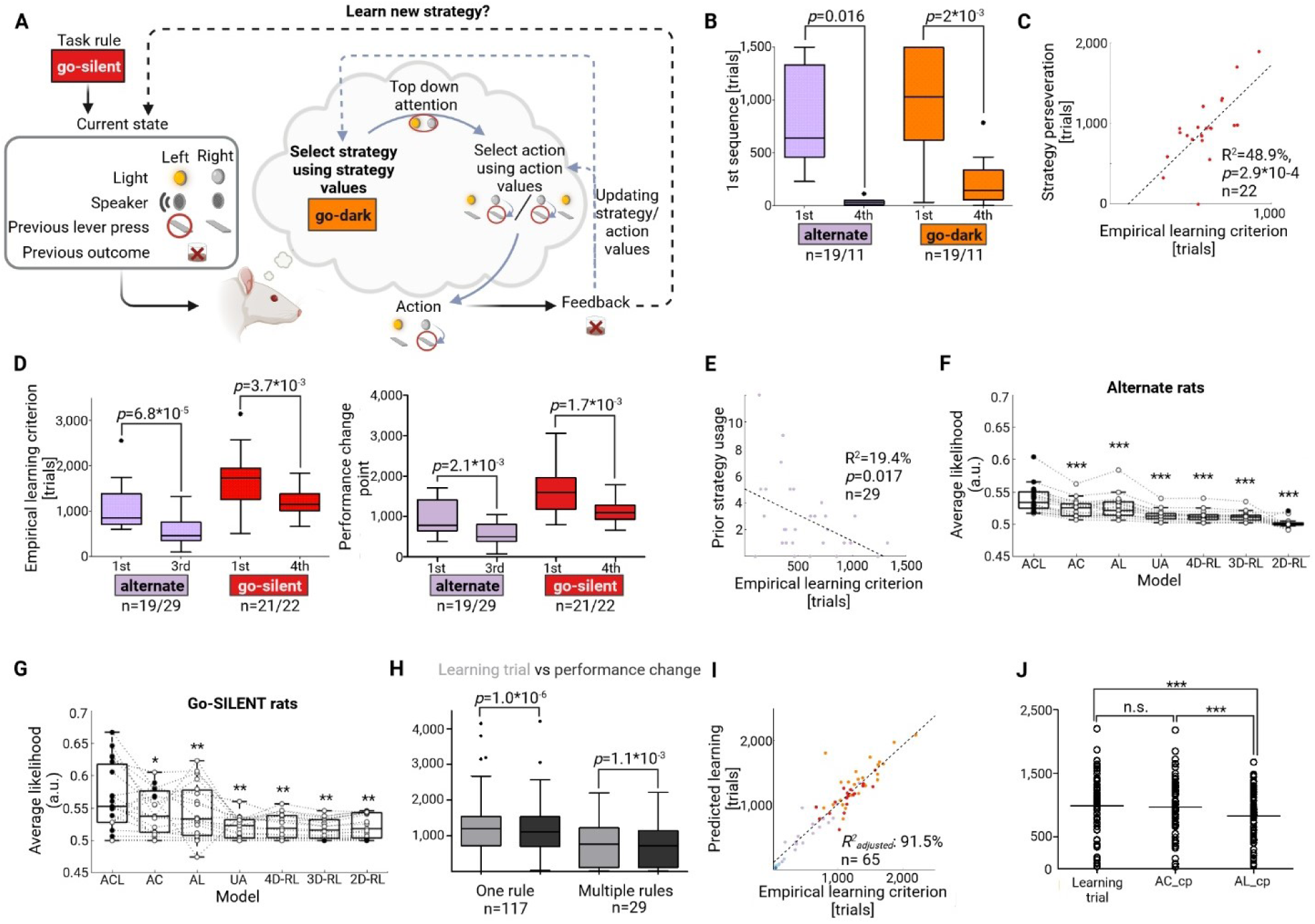
Formation and value-based selection of behavioral strategies in rats. (A) Hierarchical RL models predict that strategies are selected based on their value and need to be formed if the strategy that matches the current task rule is not part of the behavioral repertoire. (B) In the absence of a task rule, strategies are selected earlier if they have been previously reinforced (i.e., in experienced rats/fourth rule) as compared to a naïve cohort (first rule), Mann-Whitney test. (C) Perseveration on the previously reinforced strategy *alternate* predicts when the rule go-silent is learned, indicating a negative transfer effect (each dot corresponds to one rat). (D) Hierarchical models also predict positive transfer effects. Indeed, both the learning trial (*left*) and the performance change point (*right*) occurred earlier in experienced rats learning the task rules alternate/go-silent as compared to naïve animals (Mann-Whitney test). (E) The number of times a rat used the strategy *alternate* in previous rules predicted how fast a rat will learn the task rule alternate. (F, G) Replication of RL model findings for rats learning alternate (F) and go-silent (G) as a first rule. Whiskers: 1.5 IQR; dashed lines connect values (circles) of the same rat; filled circles represent best (i.e., highest likelihood) model of each rat. Model comparisons between ACL and all other models are Benjamini-Hochberg corrected Wilcoxon matched pairs test on non-normalized BICs with with *: p<0.05, **: p<0.01, ***: p<10^-3^. (H) Performance change points precede the learning trial both in rats learning one (n=117) or multiple rules (n=29), Wilcoxon signed-rank test. (I) Abrupt changes in attention (during choice formation and reward feedback) across trials predict learning. Each dot represents the empirical vs. predicted learning trial. Rules are color-coded as in Figure 1. See also Table S3, Figure S5. (J) Dot plots (horizontal line indicates group mean) show that attention-at-reward change points (AL_cp) preceded both attention-at-choice change points (AC_cp) and the learning trial (repeated-measures ANOVA, p<10^-4^, F=18.7, Bonferroni’s multiple comparison test with ***: p<10^-3^ for significant pairs).

At the behavioral level, two core predictions are that new strategies will be formed if the experimental rule is not part of the behavioral repertoire and that established strategies are selected based on their acquired value (i.e., action values at the higher hierarchical level; Figure 3A). We first tested the assumption of value-based strategy selection and then evaluated the evidence for strategy formation.

According to our hierarchical strategy selection hypothesis, previously reinforced strategies have a higher value and should thus be chosen earlier in a novel context. We therefore compared strategy selection between two cohorts of rats receiving random reward feedback that were either naïve or that previously learned three different experimenter-defined rules (go-dark→ place→ alternate). Indeed, the strategies *go-dark* and *alternate* were chosen earlier in the cohort with prior learning experience (Figure 3B). Value-based strategy selection should also go along with both negative and positive transfer effects after prior learning because strategies are re-selected as a whole instead of re-learning all possible state-action mappings.^3^ Indeed, we observed that the learning trial for the rule go-silent after a sequence of three experimenter-defined rules (go-dark→ place→ alternate) was delayed depending on how much rats were perseverating on the strategy *alternate* (Methods, Figure 3C). However, overall positive transfer effects were observed such that experienced rats learned the task rules alternate and go-silent (as a third and fourth rule) faster than naïve animals (Figure 3D). The number of times a rat used the strategy *alternate* in previous task rules (which indicates a higher strategy value) also explained some of the learning variance for the task rule alternate (Figure 3E). Further, RL model-based analysis revealed that the behavior of naïve rats acquiring the rules alternate or go-silent as a first rule is best explained by the ACL model (Figures 3F, 3G), suggesting that slower learning of the first rule is not caused by a qualitatively different learning process. This being said, we observed that many rats did not use the correct strategy before it was required (Figure 3E). This indicates that positive transfer effects cannot be entirely explained by differences in acquired strategy value but may also be caused by how fast rats add a new strategy to their repertoire.

We reasoned that the formation of a behavioral strategy requires that rats sample the relevant state-action space first to learn correct mappings at the low level. This will increasingly affect action selection and eventually a novel high-level abstract strategy is formed. To test this assumption, we first tried to identify behavioral markers related to action selection that predict the learning trial. Indeed, the performance change point does not only correlate with learning (Figure 1E) but also precedes the empirical learning criterion both in rats learning one rule and multiple rules (Figure 3H). Moreover, we confirmed that the performance change point occurs earlier in experienced vs. naïve rats learning the task rules alternate and go-silent (Figure 3D). We also used change point analysis to test if attention increases during rule acquisition and whether such transitions are predictive of learning. While attention-at-choice in RL models reflects trial-by-trial fluctuations when rats focus their attention on different task features, this analysis focused on how attention for the strategy-specific task feature changes during rule learning. Indeed, we detected transitions in attention measures in a large proportion of learned rules (attention-at-choice with respect to the strategy-specific task feature: 88.1%, attention-at-reward: 66.1%; Figure S5, Table S3). In line with the idea that strategy formation requires increased learning, the first correct strategy sequence occurred much later in rules for which an attention-at-reward change point could be detected (trial 414 (182.3-817.8), n=72 rules) as compared to rules without such a change point (trial 53 (34-257), n=37 rules; p=9*10^-6^, Mann-Whitney test). We then estimated a linear regression model to show that both attentional change points (attention-at-choice p=1.9*10^-15^, attention-at-reward p=7.3*10^-4^) predict the learning trial with high accuracy (R^2^_adjusted_=91.5%; Figure 3I, Table S3). Change points for attention-at-reward preceded both attention-at-choice change points and the learning trial but the latter two were not different (Figure 3J).

In sum, rats learn new strategies if the behavioral repertoire does not match current task demands. Moreover, value-based selection of established strategies is an important action selection mechanism during multidimensional rule learning that explains transfer effects observed in the data.

### Abstract representation of strategies and task features in rat PFC

Research on hierarchical rule learning in humans also suggests that strategies as abstract, hypothetical rules are represented in prefrontal brain activity.^3^ Using multiple single-unit recordings, we found that behavioral strategies could be decoded from the activity of prelimbic neurons in rats performing rule switches in either the novel multidimensional rule-learning paradigm or a conventional set-shifting task (62 sessions from nine rats, 34 (21-38) neurons per session; Methods, Figure 4A). Maximum decoding accuracy from population activity for each pair of strategies (n=105) within an experimental session was 80.9 ± 1.0% and peaked at 0.66s (-1.05 - +1.2) after lever onset. Interestingly, decoding accuracy was already above chance three seconds *prior* to cue onset (72.5 ± 1.2%, Figure 4B). Electrophysiological findings thus corroborate behavioral results, which indicate that action selection is a top-down process and not tied to specific cues.

**Figure 4.**
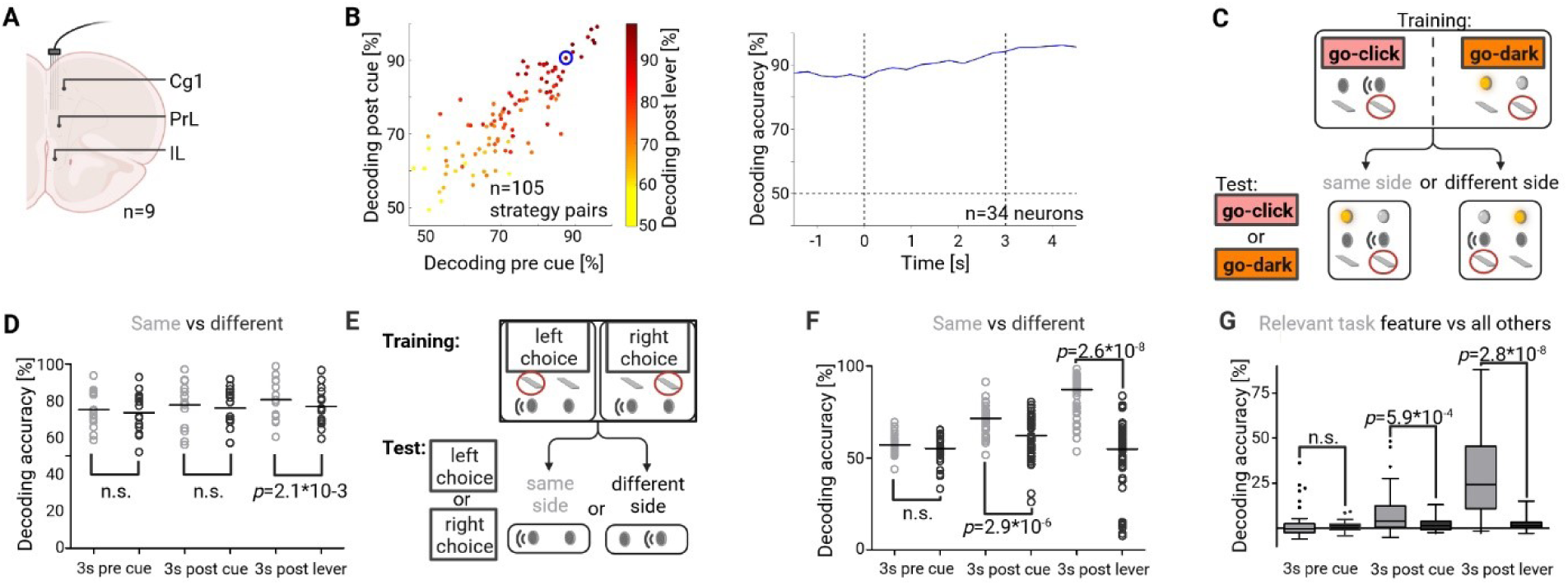
Abstract representation of strategies and task features in rat PFC. (A) Population decoding of selected strategies based on prelimbic multiple single-unit activity recordings. (B) *Left:* decoding accuracy is shown for three consecutive 3s bins of each strategy pair. Accuracy is already above chance *prior* to cue onset, indicating that action selection is a top-down process and not driven by cues. *Right:* decoding accuracy across time shown for the strategy pair highlighted with a blue circle in the left panel (dotted horizontal line at 50% corresponds to chance level, vertical lines indicate cue and lever onset). (C) Prefrontal representations of strategies did not depend on motor response in a given trial. A classifier was trained on trials with lever presses on one side and decoding accuracy was compared when tested on trials with lever presses on either the same or opposite side (example shown *left*). (D) Decoding accuracy was not different between conditions in the 3s before and after cue onset but significantly lower in the 3s following cue onset. (n=14, dot plots with mean as horizontal line, paired t-test). (E) Analysis strategy to test whether action decoding depends on a specific task feature (example sound). (F) Action decoding depended on the relevant task feature, most prominently following lever onset (n=41, dot plot with median as horizontal line, Wilcoxon matched pairs test). (G) The difference in decoding accuracy between same and different was plotted for three consecutive time bins both for the rule-relevant task feature and the median of all other task features. Decoding asymmetry was task feature-specific for both the 3s post cue and lever onset (n=41, whiskers: 1.5 IQR, Wilcoxon matched pairs test).

Moreover, hierarchical RL models also predict that strategies should be abstract “high-level” actions (Figure 3A), which implies that the prefrontal representation of strategies should be invariant to “low-level” motor responses in a given trial.^34^ To test this prediction, we trained a classifier to discriminate between strategy pairs, but only using trials where all lever presses were on one side (i.e., all trials with either left or right lever presses for both strategies). We then compared strategy decoding in two testing conditions: either using trials where all lever presses were on the same side used for training, or on the opposite side (Methods, Figure 4C). Our results for three consecutive time bins within a trial confirmed this prediction: decoding accuracy did not decrease in the opposite side condition compared to the same side condition in the three seconds prior or following cue onset. However, there was a significant difference for the three seconds following lever onset (Figure 4D). These findings show that the chosen lever can be decoded from prefrontal population activity, but only at the time of lever press, indicating that this representation is not driving low-level action selection. Instead, our findings demonstrate that the prefrontal cortex rather represents high-level behavioral strategies and that this representation has features of an abstract rule, not tied to simple motor actions.

We used a similar analysis approach to test whether there was electrophysiological evidence that rats have a representation of the relevant task feature even before they infer the correct task rule (i.e., while they are sequentially testing different hypotheses). We decoded whether rats pressed the right or the left lever and evaluated whether decoding accuracy depended on the task feature that is relevant for the current task rule (Methods). More specifically, in the example given in Figure 4E, all right and left choice trials of a session (i.e., no matter whether trials were assigned to a specific strategy or not) were sorted according to whether the right or left loudspeaker was active in a given trial. We then trained the decoder on left-right choice pairs for which the loudspeaker was active on the same side and tested on pairs with the active loudspeaker on either the same or the opposite side. We reasoned that action decoding is dependent on a task feature if decoding accuracy is significantly higher in the condition where the side of the active loudspeaker is the same in both the train and test condition. We found that action decoding depended on the relevant task feature (e.g., sound for rats learning the rule go-silent) in the three seconds following cue and lever onset (Figure 4F).

Furthermore, this decoding asymmetry (i.e., the difference in decoding accuracy between same vs. different) was specific for the task-relevant feature. We showed this in two different ways. First, we compared the difference between the decoding accuracy for the relevant task feature compared to the median of all other task features. We did not detect a significant difference for the three seconds prior to cue onset. However, we found higher decoding asymmetry for the relevant task feature during cue presentation and after lever onset (Figure 4G). Second, we counted how often the decoding difference was >5% for the relevant task feature vs. all other features. Again, there was no difference for the three seconds prior to cue onset (7/41 vs. 16/115 sessions with >5% difference, p=0.62, chi-square test) but for the three seconds post cue (19/41 vs. 27/115, p=5.8*10^-3^) and post lever onset (38/41 vs. 22/115, p=1.1*10^-16^).

In sum, we detected an abstract neural representation of specific strategies throughout task stages, which is in line with their proposed role in top-down action selection. Later in a trial, prefrontal activity also reflected the relevant task feature independent of the current action (i.e., this effect was not restricted to specific trials or sessions within a rule).

### Strategy-based learning in humans

Although our rodent task is inspired by concepts from human research, it is unknown whether humans show similar behavioral and neural processes in this specific experimental context. We therefore analyzed data from 31 healthy adults that solved an adapted version of the multidimensional rule-learning task during MEG recordings (Figure 5A). Due to limits on MEG session duration, the number of trials was necessarily lower in the human task version. Nevertheless, subjects tested most of the behavioral strategies (6 (5-6) out of 8 strategies) and were able to learn 58.1% of the rules. Across rules, the onset of the first correct strategy sequence correlated with both increases in performance (Figure 5B) and decreases in reaction time (Figure 5C). Interestingly, the onset of the first correct strategy sequence occurred before the decrease in reaction time (trial 29 (15-63) vs. 51 (40.5-86.5), p=1.2*10^-3^, Wilcoxon matched pairs test). Faster responses, indicating higher confidence in their choices thus occurred only after subjects tested the correct strategy for the first time. This supports the notion that humans indeed test strategies to identify the correct task rule.

**Figure 5.**
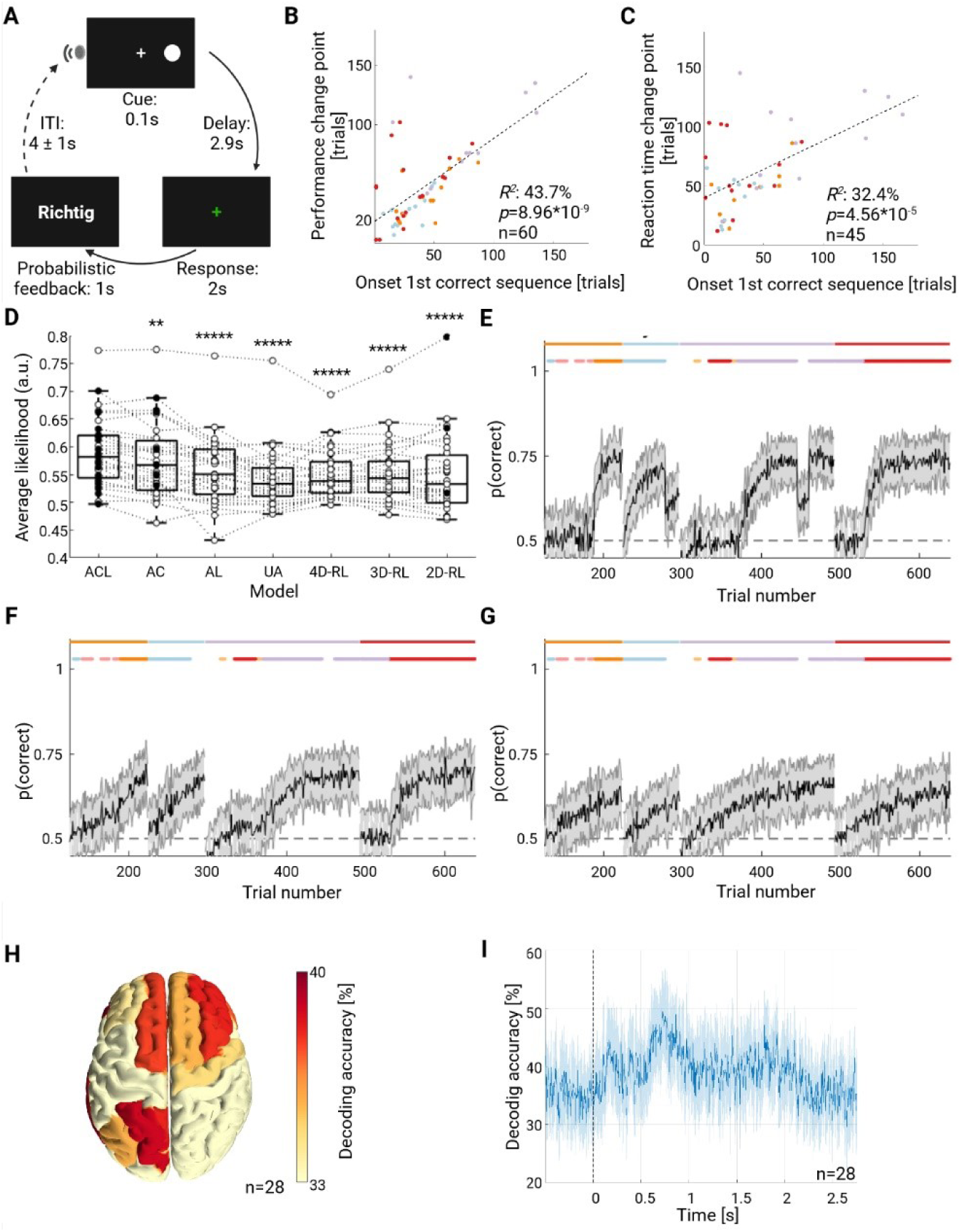
Behavioral and MEG evidence for behavioral strategies in humans. (A) Healthy adults solved an adapted version of the multidimensional rule-learning task during MEG recordings. (B) Significant correlation between the onset of the first correct strategy sequence and the performance change point mirrors rat findings. (C) Significant correlation between the onset of the first correct strategy sequence and a decrease in reaction time. (D) Replication of RL model findings in humans. Whiskers: 1.5 IQR; dashed lines connect values (circles) of the same subject; filled circles represent best (i.e., highest likelihood) model of each subject. Model comparisons between ACL and all other models are Benjamini-Hochberg corrected Wilcoxon matched pairs test on non-normalized BICs with **: p<0.01, ****: p<10^-4^, *****: p<10^-5^. (E-G) Representative example of model simulations using model parameters estimated from the experimental data of one human. Step changes in performance could only be reproduced if choice was modulated by the strategy-specific attention focus (ACL in E), but not AL (F) or 4D-RL models (G). (H) Decoding of the task feature tested for relevance by a strategy shown at the source level over the averaged time window of 0 to 1 second after stimulus onset using a searchlight approach (33% corresponds to chance level). The spatial topography of decoding accuracy reveals a predominantly prefronto-parietal network, including superior and middle frontal gyrus, cingulate, superior parietal gyrus and precuneus. (I) Trial-based decoding accuracy shown across time with respect to cue onset (vertical dashed line), 33% corresponds to decoding at chance level. Task rules and strategies are color-coded as in Figure 1.

We investigated evidence for strategy-based learning further by fitting our attention-modulated and standard RL models to human behavior. While we lacked an empirical measure of attention in the human cohort, we used the output of our strategy detection algorithm as a binary indicator of attention to a specific strategy, and hence, a specific task feature. As in the rat cohort, we found that the ACL model was the best at predicting held-out behavioral data in 19 out of 31 subjects, followed closely by the AC model in 9 out of 31 subjects (Figure 5D). Again, model simulations constrained by human experimental data showed that experimentally observed step changes in performance could only be reproduced if choice was modulated by the strategy-specific attention focus (Figure 5E-G).

At the neural level, we decoded the task feature that is tested for relevance by a strategy instead of directly decoding strategies as in rats. This was necessary for the (random-effects) group-level analysis because subjects did not all use the same strategies but sampled the same task features (e.g., *go-click/go-silent* share the same task feature). We used multivariate pattern searchlight analysis at the source level (Methods) and found significantly above-chance decoding accuracy in a cortical network prominently including lateral and medial prefrontal as well as cingulate sources (Figure 5H). As for animals, decoding accuracy was already above chance prior to cue onset (Figure 5I), indicating that attention allocation in humans is also driven by top-down processes related to strategy learning rather than being tied to specific cues. In sum, behavioral and neural findings in humans replicate those in rodents, suggesting that both species address novel task challenges by probing different low-dimensional behavioral strategies, in contrast to solving the much higher-dimensional problem of learning all possible cue-action pairs.

## Discussion

Identifying the computational processes of executive functions and their neural implementation^36,37^ is crucial to gain a mechanistic understanding of cognitive flexibility and how it is impaired in psychiatric conditions.^1,38^ However, a key challenge on that path is the explanatory gap between macroscopic human brain imaging and the microcircuit level in animals.^36,39^ A cross-species approach has been proposed to overcome this barrier that includes aligned task paradigms in both species with detailed computational modelling of cognitive processes and the use of complementary neural activity measures that can be compared in a common representational space.^36,39–41^ However, careful behavioral work is central to interpreting neural data:^42,43^ neural representations are only informative if we can make sure that cross-behavioral assays engage similar cognitive processes. This is important because most learning problems can in principle be handled by many different computational mechanisms^37^ and humans have remarkable abilities to learn in complex environments^16^ that may be qualitatively different from how animals solve the same task.^44^ Therefore, a key aspect of our study is that it goes beyond performance measures used in traditional rule learning paradigms^1^ and evaluates whether there is a conserved set of computational mechanisms at work when both species are challenged in a matched paradigm. We used a combination of behavioral, computational and electrophysiological methods at the single-trial level to show that both rats and humans sequentially follow different hypothetical rules - which we call *behavioral strategies* - to identify the current task rule. The use of low-dimensional behavioral strategies to test task features for relevance contrasts with standard RL models that predict unstructured learning of all possible mappings between environmental states and actions.

Several computational models can explain how humans infer task rules in complex environments.^3,5,16,27^ In rodents, however, these learning mechanisms are largely unknown in spite of a wealth of information about task-related neural firing patterns, necessary brain regions and involved neurotransmitters in the context of rule-switching paradigms.^33,38^ We therefore designed rat experiments to test behavioral and neural predictions made by different learning models and then evaluated whether these computational processes are at work in a matched paradigm in humans.

An important class of models suggests that attention mechanisms can reduce the perceived complexity of a multidimensional environment by focusing the animal’s cognitive resources on a small subset of features, thus allowing for efficient sampling of a simplified state space.^11^ In line with previous literature^45^, our video analysis showed head movements related to cue onset and reward feedback and these orientation reactions were strongly modulated by strategy-specific attention. Attention-modulated RL models confirmed that task responses were best explained if empirical measures for attention at both choice and learning were included in the model, similar to what has recently been described in humans.^11^ Furthermore, simulations showed that only models that include an attention-at-choice component were able to reproduce step changes in performance data. Interestingly, similar attentional mechanisms also underlie the flexibility and generalization capabilities in powerful state-of-the-art AI systems based on Transformers.^46–48^

A challenge for attention-modulated RL models is to predict how both a strategy and the associated attention focus are selected.^5,11^ Using behavioral and neural analyses, we provided evidence that hierarchical RL, an influential model of human cognitive flexibility,^3,27^ may also account for strategy learning in rats. At the behavioral level, we confirmed that new behavioral strategies are added to the repertoire based on task demands and that established strategies are selected based on their acquired value (i.e., previously reinforced strategies are preferably selected in a novel context). Future data-guided computational modelling of high-level, strategy-based learning could complement our current low-level attention-modulated RL approach to provide novel, testable predictions on how subjects take advantage of regularities in the environment to generalize established behaviors to new situations.^5,22^

At the neural level, we decoded strategy-related quantities from prefrontal network activity of both species. Our human findings are consistent with the literature that proposes that hypothetical rules are represented by lateral PFC.^11,35^ A translational link between neural measures is challenging because large parts of the human PFC probably have no clear corresponding homologues in rodents.^49^ However, individual prefrontal neurons in both rodents and primates have consistently been found to encode abstract task rules in the steady state (i.e., *after* the task rule has been reliably identified), indicating that at least some neural computations are conserved.^13,50^ Moreover, recent work indicates that meaningful translational links for higher cognitive processes can be made even if neural recordings have different recording scales and modalities: similar neural representations in humans and mice could be identified using a common representational space that was based on a same data-analytical framework.^36,40^ Our results on strategy decoding resonate with that finding and highlight the potential of an aligned, yet complementary approach in both species. For example, higher neural signal-to-noise ratio along with slower learning curves allowed us to go beyond simple strategy decoding in rats to show how neural activity represents task structure. More specifically, prefrontal computations (of the relevant task feature or strategies as abstract actions) may support generalization because they reflect only the shared structure in the environment with unnecessary details discarded. Future work may build on these results to understand “slow” computational processes that ultimately lead to the formation of new abstract hypotheses in rats and how these are orchestrated with “fast” cognitive processes like the identification of the relevant task feature or sequential hypothesis testing.^16,51,52^ Applying such a cross-species approach is not only relevant for basic neuroscience but may also increase predictive validity in animal models of neuropsychiatric diseases.^39^ In turn, these insights could inform studies in clinical populations that combine empirical attention measures (i.e., using eye tracking), imaging and computational approaches to better understand how altered neural representations lead to impaired cognitive flexibility.^37^

## Acknowledgments

We thank the lab of Prof. Hamprecht for help with video analysis, Dr. Enkel for support during development of the paradigm and Prof. Draguhn for discussions. Some of the analyses required the use of a high-performance computing cluster (bwUniCluster 2.0, https://www.bwhpc.de/). The authors acknowledge support by the state of Baden-Württemberg through bwHPC. All figures were created using BioRender.com. Figures include color specifications and designs developed by Cynthia Brewer (http://colorbrewer.org/). This work was funded by the German Research Foundation (BA 5382/1-1, BA 5382/2-2 to F.B., DU 354/8-2, DU 354/10-1, DU 354/13-1 to D.D.), the third CIMH Young Investigator Award and the MACS (Mannheim Advanced Clinician Scientist) Program (F.B.).

## Author contributions

Conceptualization, F.B., T.P., A.M.L., H.T. and D.D.; Methodology, F.B., T.P., G.K., H.T. and D.D.; Software, F.B., T.P. and H.T.; Formal Analysis, F.B., T.P., N.B., S.H., H.Z. and H.T.; Investigation, F.B., T.P., N.B., S.H., T.M. and H.Z.; Writing – Original Draft, F.B., T.P. and H.T.; Writing – Review & Editing, F.B., G.K., A.M.L, H.T. and D.D.; Visualization, F.B., T.P., N.B. and H.T.; Supervision, F.B., T.P., A.M.L., H.T. and D.D.; Funding Acquisition, F.B. and D.D.

## Methods

### Resource Availability

#### Lead contact

Further information and requests for resources should be directed to and will be fulfilled by the lead contact, Florian Bähner.

#### Materials availability

This study did not generate new unique reagents.

#### Data and code availability

Rat and human data sets as well as code will be made available on heiDATA (https://heidata.uni-heidelberg.de/) upon publication. One human participant did not give consent to share data. Any additional information required to reanalyze the data reported in this paper is available from the lead contact upon request.

### Experimental Model and Study Participant Details

#### Animals

216 male Sprague Dawley rats (Charles River, Germany) were six weeks old when they arrived at our animal housing facility. They were group-housed in standard macrolon cages (55×33×20 cm) with free access to drinking water throughout the study. A subset of rats (n=15) was implanted with silicon probes, and these rats were single-housed after surgery in the same cage type with a custom-made lid to prevent implant damage. Food was available *ad libitum* for the first two weeks. Thereafter food was restricted to ∼20 g per rat per day which maintained them on a stable bodyweight. Lights were turned on from 7:30 am to 7:30 pm and experiments were performed during the light phase. At the start of experiments, rats were at least eight weeks old. The head and neck of rats (except for implanted rats) were marked with a black hair dye (Koleston Perfect 2/0, Wella, Paris, France) to facilitate detection of body parts in video analyses. All experiments in this study were performed in accordance with national and international ethical guidelines, conducted in compliance with the German Animal Welfare Act and approved by the local authorities (Regierungspräsidium Karlsruhe, Germany). Efforts were made to reduce the number of animals used, and all behavioral protocols were refined to minimize adverse effects on animal well-being. Throughout the study period, no adverse health events occurred that demanded special veterinary care or removal of an animal from any experiment. 17 rats were excluded from data analysis either due to incomplete data (hardware/software problems or human error, n=12) or bad signal quality in the recovery period after surgery in the case of implanted rats (n=5). Table S4 provides an overview of how the remaining 199 rats were distributed across experimental groups.

#### Human subjects

32 healthy adults without a history of mental disorder (22 females/10 males, median age 24.5, range 19-55 years) were recruited from the local community. Participants had normal or corrected-to-normal vision. All subjects gave written informed consent and received a financial compensation of 50€ for participation. The participants were invited to two appointments. The first one consisted of a magnetoencephalography (MEG) session during which participants performed the multidimensional rule-learning experiment. The second consisted of an anatomical magnetic resonance imaging (MRI) scan and a 90-minute neuropsychological test battery (CANTAB, Cambridge Cognition, Cambridge, UK) covering several cognitive domains with a focus on executive functions (order of tasks: Reaction Time/RTI, Intra-Extra Dimensional Set Shift/IED, Paired Associates Learning/PAL, Spatial Span/SPP, One Touch Stockings of Cambridge/OTS, Stop Signal Task/SST, Rapid Visual Information Processing/RVP, Spatial Working Memory/SWM). Data from the second appointment were not used for this study. One subject discontinued participation in the study prematurely; three subjects could not be included in the MEG analysis (two due to strong artifacts, one subject for behavioral reasons that are detailed in the section on MEG analysis). The study was approved by the local ethics committee.

### Method Details

#### Rat operant procedures

Initial operant training took place in automated operant training chambers (20.5×24.1cm floor area, 29.2cm high; MED Associates, St. Albans, VT, USA) that were equipped with two retractable levers, located left and right from a central food tray. Cue lights were located above each lever and a house light was placed in the upper left corner opposite the food tray. All chambers were light-and sound-attenuated and a ventilator provided constant background noise. All procedures were controlled by a computer running custom-made MedStat notation code (MedPC IV, MED Associates, St. Albans, VT, USA). Initially, rats were trained to respond equally to the presentation of both levers individually.^33^ Rats were exposed to the experimental stimuli before the actual experiment started (i.e., sensory stimuli were not novel). The rule-learning tasks were performed either in the same boxes (operant two-rule set shifting task with deterministic reward feedback) or in large (30×48×41cm, custom-made) operant boxes (all rats performing the multidimensional rule-learning task and implanted rats performing an operant two-rule set shifting task with probabilistic reward feedback). For the multidimensional rule-learning task, operant chambers were additionally equipped with two loudspeakers located above each lever. We used either sweetened condensed milk (small boxes: 80μl, Milchmaedchen, Nestlé, Germany) or food pellets (big boxes: 45mg food pellets, BioServ F0021, Flemington, NJ, USA) as rewards.

#### Rat multidimensional rule-learning task

A pseudorandomized list was used for presenting one visual and one auditory cue (click sound) at the beginning of each trial (Table S5). Three seconds after trial onset, two levers were presented. Rats had to respond within 10s after presentation, otherwise the trial was considered an omission trial. At the end of the trial, the levers were retracted and the next trial was started after a fixed inter-trial interval (ITI) of 20s. Each session consisted of 300 trials (i.e., independent of performance) and no rule switches occurred within a session. Nine different experimenter-defined rules (go-light, go-dark, go-click, go-silent, alternate, go-right, go-left, win-stay-lose-shift, random) were used. The random rule refers to a session with random reward feedback. Rats learned either a single rule or one of two versions of a fixed sequence of four rules. The go-left rule was excluded for rats learning only one rule. The sequence go-dark→ place→ alternate→ random (n=11) was used to test the effect of prior learning on strategy selection in the last rule. Five random rule sessions were performed and compared to the first five sessions of a separate cohort of naïve rats (n=19) also receiving random reward feedback. The sequence go-dark→ place→ alternate→ go-silent (n=22) was used to investigate positive transfer effects (e.g., accelerated learning of the go-silent rule as the fourth versus the first rule). In both four-rule task versions, the *place* strategy (i.e., always pressing the right or the left lever) that rats spontaneously used less during learning of the go-dark rule was selected as the second experimenter-defined rule for each rat (go-right: n=22; go-left: n=11). In multi-rule experiments, un-cued rule switches occurred after the percentage of trials explained by the correct strategy (i.e., the one corresponding to the task rule) exceeded 50% of all trials in the session or if a rat reached the criterion of 18 correct trials out of 20 in two consecutive sessions. Reward feedback was deterministic except for the random rule (50% reward independent of choice), and win-stay-lose-shift (80% reward for correct responses, 20% reward for incorrect responses; otherwise, it would not be possible to tell whether a rat followed *win-stay-lose-shift* or a *place* strategy).

#### Rat conventional set-shifting paradigm

In a paradigm adapted from Floresco et al.,^33^ the trial structure was as in the multidimensional rule-learning task. Rats first acquired a go-light rule (i.e., *“press lever with illuminated cue light above”*) until a performance criterion was reached (26/30 correct). Subsequently they switched back-and-forth between go-light and a place rule (*“always press the lever on one side”*). All rule switches were un-cued. A rat was considered to follow a given rule (i.e., having formed an attentional set) if it scored 18 correct trials out of 20. We used two versions of the task that differed with respect to reward feedback: reward feedback was either deterministic as in the original task version (two rule switches per session; min 30, max 200 trials per rule) or probabilistic (80/20% reward for correct/incorrect responses; one rule switch per session; min 30, max 250 trials per rule). The probabilistic version was introduced to encourage explorative choices and only used in implanted animals.

#### Rat surgery

15 rats underwent surgery for microelectrode implantation after they had learned to respond equally to the presentation of both levers individually. 64-channel silicon probes (chronic P1-probe with four shanks and 16 channels/shank; Cambridge NeuroTech, Cambridge, UK) attached to a drive (nano-Drive; Cambridge NeuroTech, Cambridge, UK) were implanted into the right prelimbic cortex (center of probe placed at following coordinates: AP +3.0, ML +0.6, DV -3 mm from brain surface) of rats anesthetized with isoflurane (2.0-2.5%). A bone screw above the cerebellum served as ground. Electrodes were only moved if signal quality declined. For each rat, placement of electrodes within the prelimbic cortex was confirmed using histological methods. More specifically, animals were deeply anesthetized and transcardially perfused with a 4% buffered formalin solution. The entire head with electrodes in place was kept in formalin solution for 3 weeks before brains were collected and then sectioned using a vibratome. This procedure ensured that electrode tracks were visible without further staining.

#### Rat electrophysiological recordings

Rats were allowed to recover at least seven days after surgery before they were accustomed to the recoding set-up and performed additional training sessions. The actual rule-learning paradigms started at least 14 days after electrode implantation. Multiple single-units were simultaneously recorded using a 64-channel RHD2164 amplifier connected to a RHD2000 USB interface board (Intan Technologies LLC, CA, USA). Channels were digitized with 16-bit resolution, sampled at 30 kHz and band-pass filtered between 0.1Hz and 7500Hz. The time stamps for cue lights, lever presentation and lever presses were transmitted from the Med Associates behavioral control system to the Intan recording system to align behavioral markers with neural activity.

#### Human rule switching

We developed a version of the rat multidimensional rule-learning task during MEG recordings using Presentation (Version 20.1, Neurobehavioral Systems, Berkeley, CA, USA). We aimed for a task that is both close to the rat multidimensional rule-learning task (i.e., same stimulus material, sequence of multiple, un-cued rule switches) and can be completed in a single MEG session. This came at the cost of having fewer trials as compared to the rat data and this task version was thus not designed to study behavioral parameters predicting learning. The experiment comprised 650 trials and consisted of the following five different experimenter-defined rules: random (126 trials)→ go-dark (100 trials)→ go-right (75 trials)→ alternate (199 trials)→ go-silent (150 trials). In contrast to the rodent version, reward feedback was probabilistic (80% for correct, 20% for incorrect responses) to make the task more challenging. Moreover, rule switches were independent of performance (i.e., subjects on average did not identify all experimenter-defined rules) and the length of different rule phases was based on a behavioral pilot outside the scanner (striking a balance between level of difficulty and being able to perform several rule switches in one session). The trial structure was as follows (Figure 5A): each trial started with the pseudorandomized presentation of a white circle and a sound on either the right or left side. The stimuli were presented for 0.1s, followed by a 2.9s delay interval with a white central crosshair. Participant responses (either right or left button press) were logged while the crosshair turned green for 2s. Feedback then appeared on the screen for 1s - either the word *“richtig”* (correct) as positive feedback or a white central crosshair as negative feedback. The white central crosshair was then displayed for 4 ± 1s before the next trial commenced. Before the experiment, subjects were informed about the trial structure and told that their goal was to get as many correct answers as possible. Afterwards, participants were asked whether they identified the experimenter-defined rules and how they solved the task.

#### Human MEG acquisition

MEG was recorded at a sampling rate of 1000Hz using a 306-sensor TRIUX MEGIN system (MEGIN, Finland) in a magnetically shielded room (hardware filtering: 0.1-330Hz). Signals were acquired by 102 magnetometers and 204 orthogonal planar gradiometers at 102 different scalp positions. A signal space separation algorithm implemented in the Maxfilter program provided by the manufacturer was used to remove external noise (e.g., 16.6Hz train power supply and 50Hz power line noise) and to align the data to a common standard head position across acquisition sessions based on the measured head position at the beginning of each session. Each participant’s head shape and fiducials (nasion and pre-auricular points) were digitized using a Polhemus Fastrak digitizer (Polhemus, Vermont, USA). Continuous tracking of head position relative to the MEG sensors was achieved utilizing five head position indicator (HPI) coils. Oculomotor events such as blinks and saccades were recorded using conventional vertical and horizontal electrooculography/EOG. EOG electrode impedance was kept below 10kΩ.

### Quantification and Statistical Analysis

#### Change point detection

In order to visualize changes of behavioral parameters, we used so-called CUSUM (cumulative sum of differences to the mean) plots.^53^ Formally, given a binary time series *r*_*t*_describing whether the animal was rewarded (*r*_*t*_ = 1) or not rewarded (*r*_*t*_ = 0) in trial *t*, the CUSUM performance plot is given by CUSUM(*r*_*t*_) = ∑_*t*′≤*t*_(*r*_*t*′_ − ⟨*r*⟩) where ⟨*r*⟩ is the mean of the entire time series. Candidate change points are located at extrema of the resulting curve. As an example, Figure S1 visualizes the performance change point of a rat, i.e., it does not show the performance in each trial but how it changes with respect to the mean performance across all trials. We used two different methods for statistical change point detection. For very short behavioral time series (rat two-rule deterministic set-shifting task), we used a sigmoidal learning-curve model, *x*_*t*_, based on an inhomogeneous Bernoulli process ^32^ to detect changes in performance (with *y*_*t*_ = 0 and *y*_*t*_ = 1 corresponding to incorrect and correct responses at trial *t*, respectively),

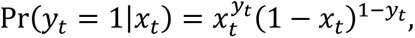

where,

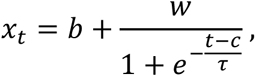

and *c* is the behavioral change point, i.e., the trial corresponding to a 50% change in correct response probability.

For all other analyses, we used the Paired Adaptive Regressors for Cumulative Sum (PARCS) algorithm. The details of the procedure can be found in^31^ and the MATLAB code is available at: https://github.com/htoutounji/PARCS. The PARCS method requires a liberal guess of the number of change points, followed by further refinement using non-parametric permutation bootstrap testing (significance level set to 0.05 based on 1000 bootstrap samples). PARCS has the advantage of detecting multiple change points in multivariate time series. However, compared to the Bernoulli approach, it is less sensitive to abrupt changes in binary, relatively short behavioral time series. In addition, we fitted a sigmoidal model to ± 100 trials centered on performance change points identified using PARCS to compute the 10-90% rise time as a measure for the abruptness of a performance change.

#### Strategy detection algorithm

For each strategy *i*, we identify trial blocks where following that strategy is more likely than chance or than following each of the seven other strategies. To do so, we first define eight binary time series ***b***_*i*_, where *b*_*ti*_ = 1 indicates that behavior at trial *t* is consistent with strategy *i*. For a candidate sequence length *n*, we compute count timeseries 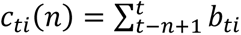, i.e., the number of trials within a window of size *n*, ending at trial *t*, for which behavior is consistent with strategy *i*, and estimate the probabilities within each of these *n*-blocks as *p*_*ti*_ = *c*_*ti*_/*n*. We then evaluate the following binomial statistic (i.e., based on the binomial distribution) at every trial,

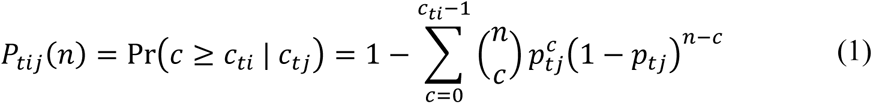

where *j* indexes the alternative strategy *i* is being tested against. When testing against chance, we set *j* = 0, *p*_*t*0_ = 0.5 and *c*_*t*0_ = *n*/2. Strategy *i* is significantly more likely followed than strategy *j* for a block of size *n* ending at trial *t* when *P*_*tij*_(*n*) < 0.05 (i.e., we ask the *H*_0_ question: “How likely is it to observe *c*_*ti*_ or more trials consistent with strategy *i* when the true strategy in place was *j*?”). Starting from *n* = 6 (the minimum admissible sequence length at which this significance level can be achieved), sequence length is iteratively increased up to the number of trials in the session. If several overlapping blocks of a candidate strategy are significant, the block with the lowest binomial statistic when compared to the second most likely strategy is kept (usually the longest) and the others are discarded. Significant blocks where the animal does not follow the tested strategy in the first or last trial in the block are discarded, as well as blocks in which the strategy is not followed for more than two trials in a row. In cases where a significant, length *m* block of one strategy *i* is a subset of a significant, length *n* > *m* block of another strategy *j*, both blocks are discarded to avoid false positives (the detection method is conservative). We excluded the strategy *win-stay-lose-shift* from strategy detection in experiments where rats learned a deterministic place rule. Empirical measures of strategy-specific selective attention were used to validate the strategy detection results (Figures S4, S5).

#### Rat empirical learning criterion

A classic performance criterion (such as 18 correct out of 20 responses) is not always a clear indication that a rat really followed a strategy consistent with the experimenter-defined rule. Increased reward rates for an incorrect strategy are possible in short blocks of trials because there are eight possible strategies but only two choices. We therefore defined an empirical learning criterion based on the length of a specific strategy sequence. We made use of the fact that strategies were also detected in an experimental cohort with random reward feedback. Strategy-specific outliers with respect to sequence length were defined as values larger than the third quartile plus 1.5 times the interquartile range (*Q*_3_ + 1.5 × (*Q*_3_ − *Q*_1_); based on 152 sessions with random reward feedback from 19 rats). We assumed that learning occurred if a detected strategy sequence length exceeded that threshold (i.e., it was an outlier). For example, for the *go-dark* strategy this threshold corresponds to a sequence length of 30 trials (Table S1). Within the few sessions an animal needs to learn the go-dark rule, the onset trial of the first *go-dark* sequence exceeding this threshold in length is defined as the “learning trial” where the empirical learning criterion is reached.

Two linear regression models were fitted to predict the “learning trial” *y*(*R*) for rule *R* which is the trial at which the empirical learning criterion was reached. The models were used to test whether strategy-related behavioral quantities are predictive of the learning trial (Tables S2, S3). To reduce the effect of potential outliers on model estimation, robust linear regression implementing the iteratively reweighted least squares algorithm (MATLAB routine *fitlm*) was used in the main text (see Tables S2, S3 for a comparison between robust and ordinary least squares results).

#### Human learning criterion

Since the task was designed to both avoid ceiling effects and to fit into one MEG session (i.e., less trials per rule), we had to adapt our criteria for learning. In our MEG sample, humans tested the correct strategy at least once in 68.5% of cases. In order to decide whether human subjects actually learned a rule, two criteria were used. Learning criterion was either reached if the percentage of trials explained by the correct strategy exceeded 50% of all trials per rule or if PARCS detected a performance change point. This allowed us to also detect learning in cases where it occurred almost instantaneously (i.e., no performance change point can be detected) or rather late with only few trials left before the next rule switch (i.e., the percentage of trials in a rule explained by the correct strategy is low).

#### Rat video analysis

A night vision USB camera (ELP, Shenzen, China) positioned above each operant chamber recorded animal behavior with 30 frames per second (Image Acquisition Toolbox, MATLAB, Natick, MA, USA). Video software (FFmpeg, https://www.ffmpeg.org/) converted the videos into grayscale, downsampled them to five frames per second and separated them into single frames. Using open source software for machine learning-based image analysis (Ilastik-1.2.0, http://ilastik.org/),^54^ frames were segmented into background, head and body of a rat. In each segmented frame, the center of head and body was determined using a custom-written Python script (https://www.python.org/). Distances between the center of the head and each of the two cue lights were determined. Additionally, the head angles with respect to both sides were computed by calculating the angles between the lines connecting head-to-body and cue light-to-body. A custom-written MATLAB script identified the frames corresponding to cue light on- and offsets in the downsampled videos. Potential markers for attention during choice formation and reward feedback were identified as follows: we observed an orientation reaction after cue onset and used the minimum angle of view with the respect to the side of subsequent responses (in the second after cue onset) as a marker for attention at choice (Figures 2, S3, S4, Table S3). Smaller angles *a*_AC_ putatively correspond to a stronger attention focus. Similarly, rats often looked back and forth between the pressed lever and the food receptacle during reward feedback. We therefore assumed that the angle sum in the second after the lever press (volatility at reward *v*_AR_) is a marker for attention-at-reward. A higher angle sum would correspond to higher attention levels.

#### Reinforcement learning (RL) models

##### Summary

We used attention-based RL models following the approach of Leong and colleagues^11^ to test whether our strategy-related attention measures are behaviorally relevant and indeed represent empirical markers for attention at choice and learning. We assumed that rats use eight strategies to probe four task features for relevance. We compared four RL models that differ in how attention modulates animal behavior: either during choice alone (attention-at-choice/AC model), learning (or reward feedback) alone (attention-at-learning/AL model), or both (ACL). This also allowed us to compare our data against the null hypothesis that attention modulates neither choice nor learning (uniform-attention/UA model). We made a further comparison between attention-based RL models and standard RL models where rats learn all possible mappings between a multidimensional state space and actions.

##### Task features, strategies and the s-a-space

At every trial *t* we assume an animal makes a choice to press a lever based on current sensory cues in addition to place and reward history. There are thus four task features that can be probed for relevance: visual (relevant for the strategies *go-light* and *go-dark*), auditory (for *go-click* and *go-silent*), place (for *go-right*, *go-left*, and *alternate*), and outcome (for *win-stay-lose-shift*). The latter relates to place strategies as well, since the decision to stay or shift depends on both place and reward history. Further, it implicitly includes the opposite strategy of *win-shift-lose-stay*, which animals rarely followed.

Formally, a trial *t* can be fully described by a combination of current and last trial features called the state ***s***_*t*_, a scalar response, or action *a*_*t*_ ∈ {*L*, *R*}, and scalar reward *r*_*t*_ ∈ {0,1}. The state is a four-dimensional binary vector ***s*** ≔ [*s*_*v*_, *s*_*a*_, *s*_*p*_, *s*_*o*_], with each entry corresponding to one of the four task features. The sensory states *s*_*v*_, *s*_*a*_ ∈ {*L*, *R*} indicate whether the visual and auditory cues in the current trial are active on the left or right side of the operant chamber. The place state *s*_*p*_ ∈ {*L*, *R*} indicates whether in the last trial the animal pressed the right or left lever. The outcome state *s*_*o*_ ∈ {*W*, *L*} indicates whether in the last trial the animal received a reward (win) or not (lose). The action *a* ∈ {*L*, *R*} indicates which lever was pressed in the current trial. An outcome-dependent action is based on the place and outcome states to indicate whether the animal pressed the same lever as in the last trial (stay), or switched to the other side (shift). As such, the eight strategies used to probe the four task features can be described by 3 state-action spaces as follows: The visual and auditory strategies are described by a 2×2 state-action space each. The place and outcome strategies are described by a 2-dimensional history-dependent state ***s***_ℎ_ ≔ [*s*_*p*_, *s*_*o*_], forming a 4×2 state-action space. More specifically, the mapping between strategies and state-action spaces is (Figure 2B): *go-click*/*go-silent* (auditory cue), *go-light*/*go-dark* (visual cue), *alternate*/*go-left*/*go-right/win-stay-lose-shift* (place/outcome).

##### Attention-modulated RL models

In these models, we assume three relevant states, the third of which is a 2-dimensional state combining the place and outcome history features. For every state *i* = 1, …, 3 and trial *t*, the animal assigns values *Q*_*ti*_(*s*_*ti*_, *a*_*t*_) to each state-action pair that reflect the animal’s expected reward from choosing action *a*_*t*_ at state *s*_*ti*_, updates these values based on reward experience, and use them to guide future choices. To estimate expected reward in a trial, we assumed the animal combined values over states linearly, weighted by attention to the respective feature state at choice time,

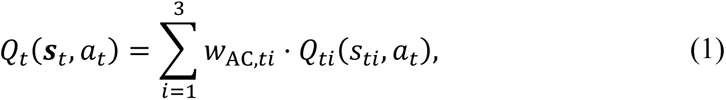

where *w*_AC,*ti*_ are attention-at-choice scores obtained by normalizing raw angle values *a*_AC_ as measured by video tracking to the range [1/3, 1] with 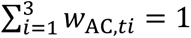 such that higher values correspond to stronger attention focus during choice.

In all models, values are updated through a Rescorla-Wagner-type learning rule,^55^

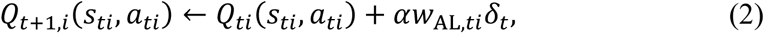

where α ∈ [0,1] is a free parameter that modulates learning rate, δ_*t*_ ≔ *r*_*t*_ − *Q*_*t*_(***s***_*t*_, ***a***_*t*_) is a scalar reward prediction error, and *w*_AL,*ti*_ are attention-at-learning scores that weigh the update for each task feature. Those scores are obtained by normalizing raw angle-sum values *v*_AR_ as measured by video tracking to the range [1/3, 1] such that higher values correspond to stronger attention focus during learning.

Given the linearly combined values over task features (Eq. 1), the policy, i.e., the probability of choosing an action given the current state, is computed using the softmax choice rule,

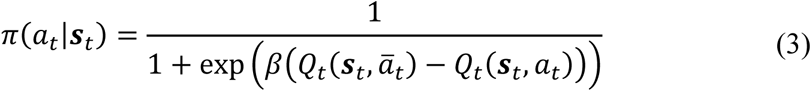

where *β* ≥ 0 is a free parameter that modulates the balance between exploration and exploitation, and *ā*_*t*_ is the opposite action of *a*_*t*_(e.g., if *a*_*t*_ = *L* then *ā*_*t*_ = *R*).

In the ACL model, learning and choice were modulated by empirically obtained attention measures ***w***_AL_ and ***w***_AC_, respectively. In the AC model, all *w*_AL,*ti*_ were set to 1/3 such that value update was uniform across all rules. In the AL model, all *w*_AC,*ti*_ were set to 1/3 such that choice in all rules was independent of attention. In the UA (uniform-attention) model, both *w*_AC,*ti*_ and *w*_AL,*ti*_were set to 1/3 such that attention has no impact on neither learning nor choice. In all models, in trials where attention was not clearly focused on a single strategy (no-attention trials), attention measures were set as in the UA model. Of note, since the attention measures were solely set according to the observed head direction angles, all models thus had the same number of free parameters (α, *β*).

In the human cohort, it was not possible to obtain empirical attention measures to different task features. However, we reasoned that our strategy-detection algorithm may indicate which task feature subjects are attending to in every trial and used binary attention measures in trials where a strategy is detected. For instance, in trials where the strategy go-silent or go-click was detected, both attention-at-choice and attention-at-learning scores for the auditory state space were set to 1 and those for the other two state spaces were set to 0. In trials where no strategy was detected, all scores were set to 1/3.

##### Standard RL models

We also consider three variations of standard RL models which directly map a multi-dimensional state to actions using 4-,3-, or 2-dimensional state spaces and that are not modulated by attention scores. In the 4-dimensional state space model, the animal assigns value *Q*_*t*_(***s***_*t*_, *a*_*t*_) to each combination of a state comprising all four relevant task features at the current trials and a binary action. This results in 32 different state-action values that are updated by a standard Rescorla-Wagner learning rule^55^,

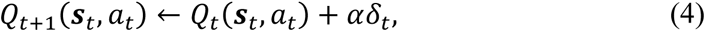

The probability of choosing an action given the current state, is computed using the softmax choice rule (Eq. 3). The 3-dimensional state space model excludes the outcome feature, resulting in 16 state-action values, while the 2-dimensional state space model excludes the place feature as well, depending only on sensory features and resulting in eight state-action values in total.

##### Model Inference, Comparison, and Simulation

In all four attention-modulated RL models, free model parameters α and *β* were inferred by maximizing data log-likelihood using MATLAB’s constrained optimization, *fmincon*, implementing the active set algorithm,

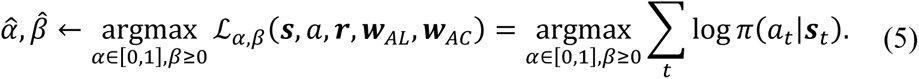

Similarly for the three standard RL models, parameters were inferred by maximizing ℒ_α,β_(***s***, *a*, ***r***). Model comparison used one-session forecasting for cross-validation. Each animal (n=29) attempted to maximize reward in *M* sequentially presented experimenter-defined rules (three or four rules with random rule sessions excluded). This required *N* sessions in total, where *N* > *M* for all animals. For each experimenter-defined rule, all sessions up to and excluding the session where animal reached criterion were used as training data (free-parameter inference with maximum likelihood). The last session per experimenter-defined rule is used for testing, where test-data likelihood is computed given the inferred parameters from the training data. This likelihood is then divided by the number of trials in the testing session to obtain trial-averaged likelihood. In total, *M* sessions were forecast per animal based on up to *N* − 1 training sessions. The resulting *M* trial-averaged test likelihoods per model per animal were then averaged to compute a goodness-of-fit score and compared between the 7 RL models (Figure 2D). A similar procedure was used to infer and compare models in human subjects (n=31, Figure 5D), and the two cohorts of animals trained on a single experimenter defined rule (go-silent: n=19, alternate: N=19, see Figures 3F, 3G).

Based on the estimated model parameters, we simulated each of the seven model variants 1000 times. To generate the simulated model learning curves and their confidence intervals (Figures 2 E-G and 5 E-G), we drew 100×100 samples of these 1000 simulations and computed the mean of each 100. This resulted in a 100 simulated mean-performance learning curves. The figures depict the median of these learning curves and the 90% confidence interval.

#### Rat electrophysiology: data preprocessing

Raw data were first bandpass-filtered between 600-6000 Hz (Butterworth filter using MATLAB function *filtfilt*) and at each time point, the median across all channels was subtracted to reduce noise and remove artifacts.^56^ Preprocessed data were then automatically spike sorted with Klusta (https://github.com/kwikteam/klusta) and afterwards manually curated with Klustaviewa (https://github.com/klusta-team/klustaviewa).^57^ During manual curation, each set of events (so-called “units”) detected by a particular template was first inspected and discarded if the events (“spikes”) comprising the unit were either judged to correspond to noise (i.e., non-physiological waveform shape or activity pattern across channels) or multi-unit activity (i.e., low-amplitude waveforms and/or multiple waveforms with refractory period contamination). The remaining units were compared to similar, spatially neighboring units to determine whether they should be merged (this was based on similarity of spike waveform/distribution across channels, drift patterns or cross-correlogram features). Similarly, a candidate unit was only kept if its spike waveforms were separated from all other units based on visual inspection of at least one 2D projection of principal component features. Finally, units were excluded if inter-spike interval violations exceeded 1% (i.e., more than 1% of consecutive spikes in accepted clusters with inter-spike interval less than 2ms). Units passing these criteria were considered to reflect the spiking activity of a neuron and included in further analyses.

#### Rat electrophysiology: decoding analyses

Decoding analyses of multiple single-unit data were performed by adapting MATLAB code from the Neural Decoding Toolbox (www.readout.info).^58^ Analyses were run using neural data from individual experimental sessions (using recordings from both the multidimensional rule-learning task and the probabilistic two-rule set shift). We included all sessions with ≥10 simultaneously recorded neurons and at least two different strategy types with ≥15 trials each (these numbers were based on toolbox recommendations^58^ and the structure of our data). Based on these criteria, 62 experimental sessions from nine rats were included in all further analyses (n=4 multidimensional rule-learning task, n=5 probabilistic conventional set shift). Raw spike trains were first aligned to trial onset (spikes from -3s to +1s with respect to cue onset were extracted) for each prefrontal neuron. Trial-aligned spike trains from trials with a detected strategy were concatenated for each neuron and labelled accordingly (i.e., each trial had a label with the specific strategy used). Spikes were then binned using a sliding window approach. We systematically tested the effect of different bin widths [250, 500, 1000, 2000, 3000ms] and respective step sizes [25, 50, 100, 200, 300ms] on decoding accuracy. We z-scored normalized the data to avoid that neurons with higher firing rates have a larger influence on the classification results. We used a maximum-correlation-coefficient^58^ classifier for decoding analyses based on leave-one-out cross-validation. The classifier learns a mean neural population vector for each class (i.e., a template for a detected strategy in our case) by averaging all training points within each class. The classifier predicts the correct class using Pearson’s correlation coefficient between the test data and the training set of each class (the highest correlation value corresponds to the predicted label). In our case, neural data from 14 trials of each strategy were used as a training set to predict which strategy a rat used in the test trial. Decoding accuracy for each session was based on 50 cross-validation runs using a resampling procedure. Accuracy on average increased monotonically [71.7 (66.4-79.6), 74.3 (66.9-83.1), 76.3 (68.9-84.7), 79.5 (69.8-88), 80 (69.1-88.3)]% and was maximal for the coarsest bin width/step size in 45/62 cases. All follow-up analyses were thus performed using the bin width with the best decoding results across sessions (3000ms bin width/300ms step size) and the decoding analysis was repeated with a bigger time window (-3s to +6s centered on cue onset) to determine the time point of maximum decoding accuracy.

All follow-up analyses were performed using the bin width with the best decoding results across sessions (3000ms bin width). We first repeated the decoding analysis with a bigger time window (-3s to +6s centered on cue onset) to determine the time point of maximum decoding accuracy. In order to make decoding accuracies comparable between sessions with different number of detected strategies (range: 2-4 per session), we performed decoding using strategy pairs we performed all further decoding analyses using strategy pairs (n=105) with the same analysis parameters.

We also conducted so-called generalization analyses which test whether decoded neural representations are abstract.^58^ This approach was used for two analyses. First, we used a subset of sessions (using pairs with at least 30 trials per strategy) to test whether the prefrontal representation of strategies (that can be interpreted as abstract actions^34^ depends on the motor response in a given trial (i.e., left versus right lever press) or is invariant with respect to that feature (Figure 4C). More specifically, we trained the classifier on the neural data (either 3s before cue onset, after cue onset or following lever onset) of 14 trials of each strategy with a motor response on one side and compared the decoding accuracy in the test set when using trials with a motor response on either the same or the opposite side (50 resample runs). Second, we decoded whether rats pressed the right or the left lever and tested whether this representation depended on a specific task feature (Figure 4F). For example, to test whether action decoding depends on the task feature auditory cue, all right and left choice trials of a session were sorted according to whether the right or left loudspeaker was active in a given trial. We then trained the decoder on left-right choice pairs (neural data either from 3s before cue onset, after cue onset or following lever onset) for which the loudspeaker was active on the same side and tested on pairs with the active loudspeaker on the same or the opposite side. We used the same analysis parameters as above (at least 15 trials for each of the four conditions, 50 resample runs). Action decoding was defined as task feature-dependent if decoding accuracy was higher in the condition where the side of the active loudspeaker is the same in both the train and test condition. All trials of a session were used with the exception of sessions in which the empirical learning criterion was reached. In this case, all trials that occurred before the learning criterion were included in the analysis.

#### Human MEG: data preprocessing

MEG data preprocessing involved the segmentation of epochs of 2s before to 4s after stimulus presentation. Eye-movement-related activity and cardiac signals were identified with independent component analysis^59^ and then discarded. All data were analyzed using the MATLAB-based toolbox for neuroelectric and neuromagnetic data analysis FieldTrip.^60^

#### Human MEG: source-level analysis

Forward models were computed based on the MNI ICBM 2009 template brain (http://www.bic.mni.mcgill.ca/ServicesAtlases/ICBM152NLin2009). A parcellation scheme based on the Desikan-Kiliani atlas^61^ was used. Following procedures similar to those described in Schoffelen et al.,^62^ single-dipole-specific spatial filters were concatenated across vertices comprising a given parcel. For each parcel, singular value decomposition was performed to extract spatially orthogonal and temporally uncorrelated components that describe the time course of activity within a parcel. These components were ordered by the amount of variance explained. Subsequently, the first principal component was selected capturing the parcel’s time course of activity. This procedure yielded 68 virtual sensors corresponding to each parcel that were subsequently treated in a similar fashion as MEG sensor level activity.

#### Human MEG: decoding analyses

Trial-based decoding was performed using the MVPA-Light toolbox (https://github.com/treder/MVPA-Light).^63^ We decided to decode the relevant task feature instead of individual strategies (e.g., *go-light* and *go-dark* share the same task feature) to be able to include all available data sets in our group-level statistics (i.e., not all subjects displayed the same behavioral strategies). Our choice was motivated by RL model findings that show that this is a reasonable assumption (e.g., *go-click/go-silent* share the same state-action pairs, Figure 2B). Since the strategy *win-stay-lose-shift* (task feature: outcome history) was not consistently detected across subjects (14/29 subjects eligible for MEG analysis), decoding analyses focused on the three remaining task features (auditory, visual, choice history; detected in 28/29 subjects eligible for MEG analysis) in 28 subjects. Classification was performed using linear discriminant analysis implementing ten-fold cross-validation with two repetitions. We computed discriminative information across time to test whether decoding accuracy depends on sensory stimuli. A searchlight approach was used to highlight the source-level topography of decoding accuracy.

#### Statistics

Data analysis was performed with Graphpad Prism (Version 7) and MATLAB (R2017a, MATLAB, Natick, MA, USA) if not otherwise noted. Normality distribution assumption was tested using the Shapiro-Wilk test. If systematic deviations from normality were detected, non-parametric statistical testing was used. Data are displayed as mean ± standard error of the mean (SEM) or as median and first and third quartile (Q_1_-Q_3_), Tukey-style whiskers were used in boxplots. Data points outside the boundary of the whiskers are plotted as outliers using individual dots. Outliers are thus defined as values greater than Q_3_ + 1.5 × interquartile range/IQR and smaller than Q_1_ − 1.5 × IQR, respectively. The significance threshold was set at p<0.05, two-sided tests were used if not otherwise specified.

## Supplemental information

**Figure S1.**
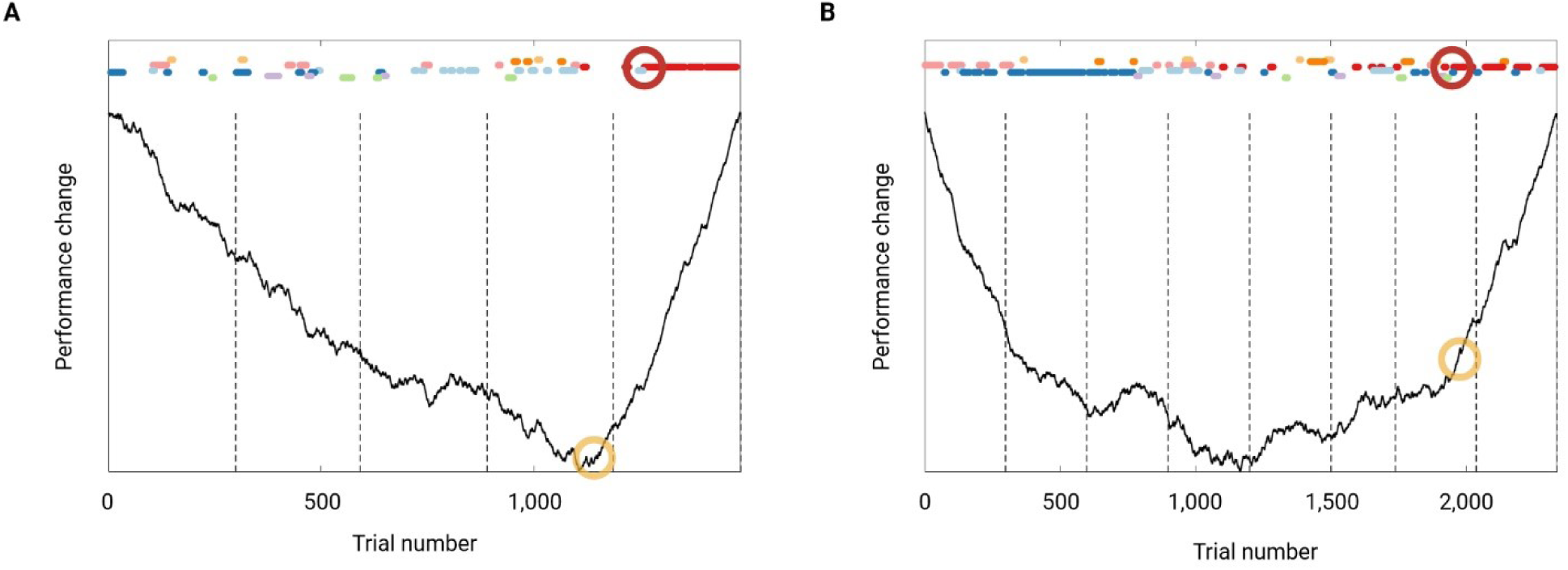
related to Figure 1. Visualization of strategy-related predictors of learning. Two examples of rats learning the rule go-silent are used to illustrate strategy-related predictors of learning (detected strategy sequences are color-coded as in Figure 1). Dashed vertical lines mark the end of an experimental session. (A) A “good learner” that reached the empirical learning criterion (which corresponds to a sequence length ≥ 21 trials) after 1260 trials (highlighted with a red circle). Change in performance is visualized using a CUSUM plot (cumulative sum of differences to the mean),^1^ a performance change point was statistically detected at trial 1139 (yellow circle). The following strategy-related response parameters were used to predict learning (empirical learning criterion) in our regression model (Figure 1F, Table S2): the trial at which the correct strategy was first tested during rule learning (here: onset of first *go-silent* sequence detected in trial 1117), a strategy-based learning rate (i.e., how many sequences occur before a rat sticks to the correct strategy; here: two *go-silent* sequences were detected before the empirical learning criterion was reached), as well as a measure for perseveration at the strategy level. This measure was chosen based on the observation that slow learners often revisited unsuccessful but salient strategies multiple times rather than testing alternative strategies for relevance. We computed how many trials rats need to “sort out” the most salient strategy and this was defined as the 10-90% rise time of the empirical cumulative distribution based on all detected sequences of that strategy. For the rule go-silent, we picked the *place* strategy the rat spontaneously used more often (Table S1). In this example, *go-right* (light blue dots) is the most salient strategy and the 10-90% rise time corresponds to 562 trials. (B) The “bad learner” needed 1955 trials to reach the empirical learning criterion and the performance change point was detected in trial 1974. The trial in which the correct strategy was first tested during rule learning was similar as compared to the good learner (trial 1100). However, eight *go-silent* sequences were detected before the empirical learning criterion was reached and strategy perseveration was 1595 trials (corresponding to the 10-90% rise time of the most salient strategy *go-left* visualized as dots in dark blue).

**Figure S2.**
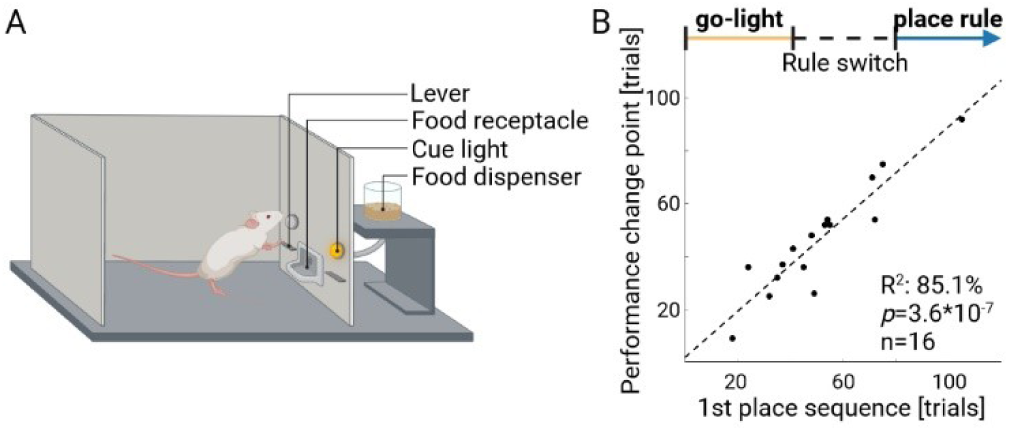
related to Figure 1. Rats also use behavioral strategies in an established two-rule set-shifting task. (A) Rats learned to switch from a go-light rule (i.e., *“press lever with illuminated cue light above”*) to a place rule (*“always press the lever on one side”*) in a classic operant set-shifting task.^2^ We also used our strategy detection algorithm and assumed that rats may use up to six different behavioral strategies in this task (*go-light/go-dark*, *go-right/go-left*, *alternate or win-stay-lose-shift*). Indeed, multiple behavioral strategies (including the strategy corresponding to the experimenter-defined rule) were detected during initial rule learning (3 (2-3) strategies tested during initial go-light rule, n=16 rats). (B) In this task, sudden transitions in performance are known to occur during a first rule switch from a go-light to a place rule.^3^ Our results indicate that these transitions can be explained by strategy switches: the scatter plot shows a significant correlation between the performance change point and the onset of the first correct *place* strategy sequence. These results are thus compatible with our hypothesis that rats follow low-dimensional behavioral strategies to infer task rules. However, it remains unclear whether the detected strategies are relevant for learning because the learning curve is rather short (performance change point occurred already after 46.3 ± 5.2 trials) and the state space of this task is very simple (two states are sufficient).

**Figure S3.**
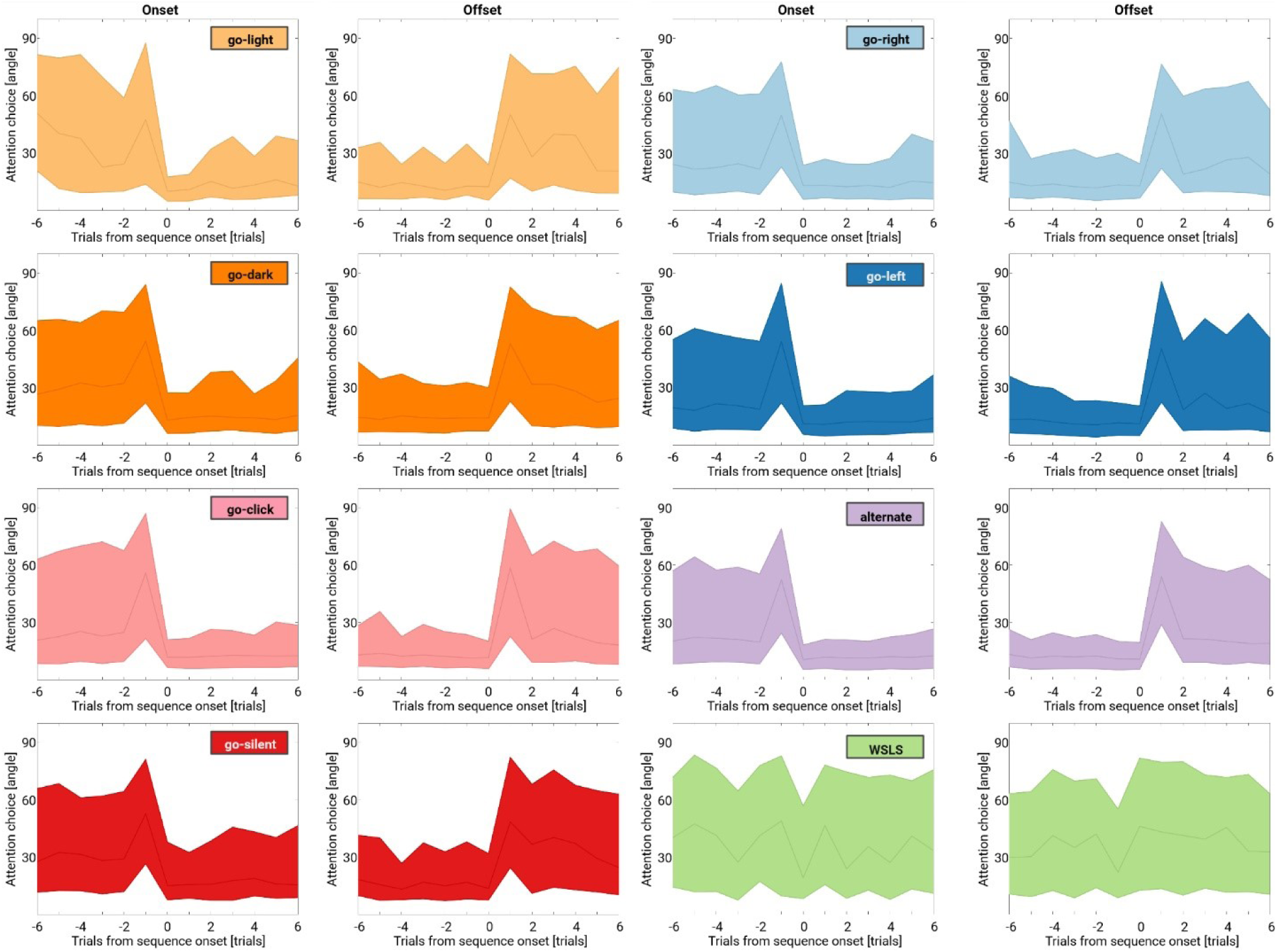
related to Figure 2. Head direction plots during choice formation for all strategies detected in the multidimensional rule-learning task. Sudden transitions in choice at sequence onset/offset were related to attention shifts for all eight detected strategies. Head-direction plots show head angles at cue onset with respect to the correct side (e.g., the side with the active cue light for the *go-light* strategy) are shown for the six trials before/after strategy onset/offset. Note that the strategy-specific head angles abruptly decreased/increased at strategy onset/offset (corresponding to trial 0). Each plot is the average of multiple strategy sequences (as detected using our strategy detection algorithm) from 29 rats performing several rule switches in the multidimensional rule-learning task. The table below lists how many sequences were detected per behavioral strategy and the p-value (Wilcoxon matched pairs test) for the decrease in head angle at strategy onset (compared to the trial before, i.e., trial -1 vs. trial 0) and the increase at strategy offset (last trial of sequence vs. trial after, i.e., trial 0 vs. trial +1).

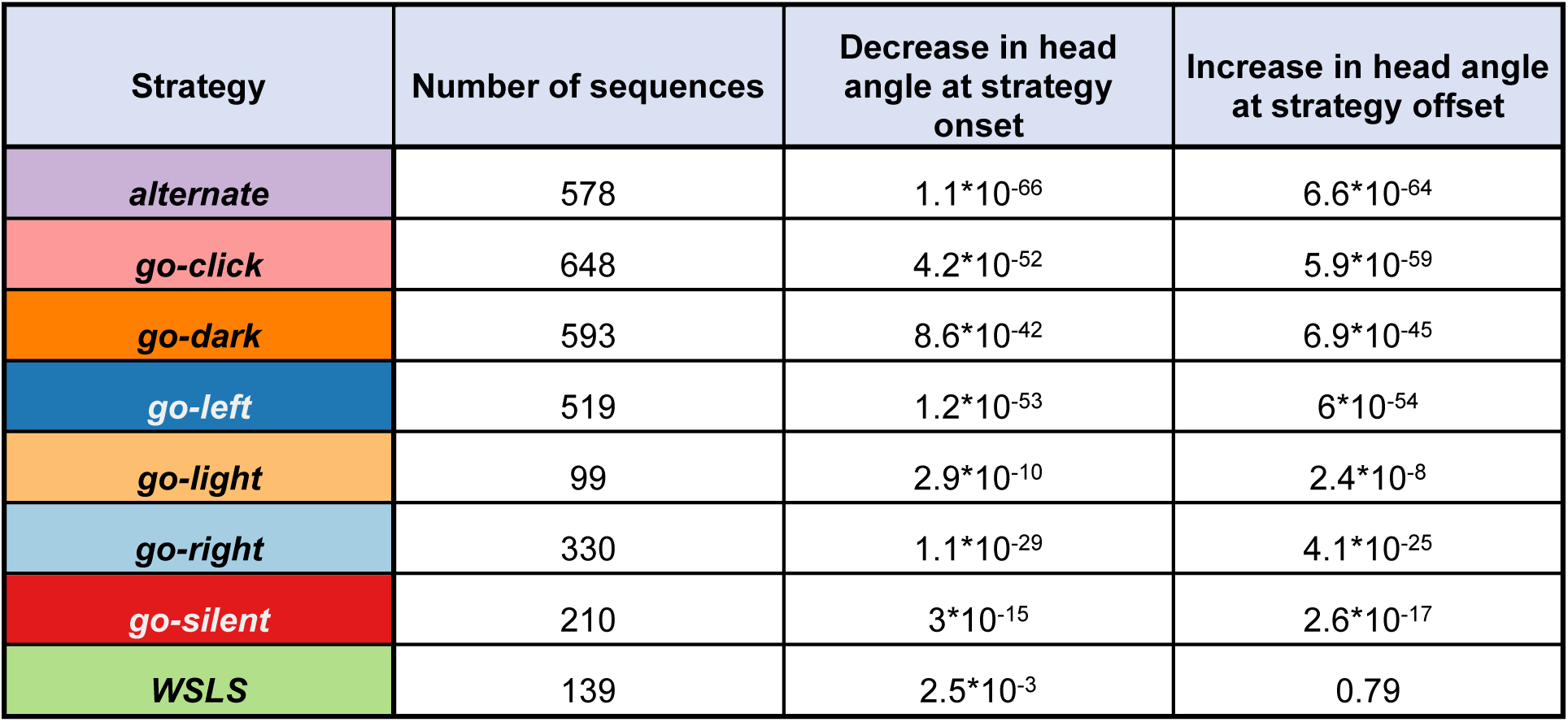

| Strategy | Number of sequences | Decrease in head angle at strategy onset | Increase in head angle at strategy offset |
| --- | --- | --- | --- |
| *alternate* | 578 | $1.1 \cdot 10^{-66}$ | $6.6 \cdot 10^{-64}$ |
| *go-click* | 648 | $4.2 \cdot 10^{-52}$ | $5.9 \cdot 10^{-59}$ |
| *go-dark* | 593 | $8.6 \cdot 10^{-42}$ | $6.9 \cdot 10^{-45}$ |
| *go-left* | 519 | $1.2 \cdot 10^{-53}$ | $6 \cdot 10^{-54}$ |
| *go-light* | 99 | $2.9 \cdot 10^{-10}$ | $2.4 \cdot 10^{-8}$ |
| *go-right* | 330 | $1.1 \cdot 10^{-29}$ | $4.1 \cdot 10^{-25}$ |
| *go-silent* | 210 | $3 \cdot 10^{-15}$ | $2.6 \cdot 10^{-17}$ |
| *WSLS* | 139 | $2.5 \cdot 10^{-3}$ | 0.79 |

**Figure S4.**
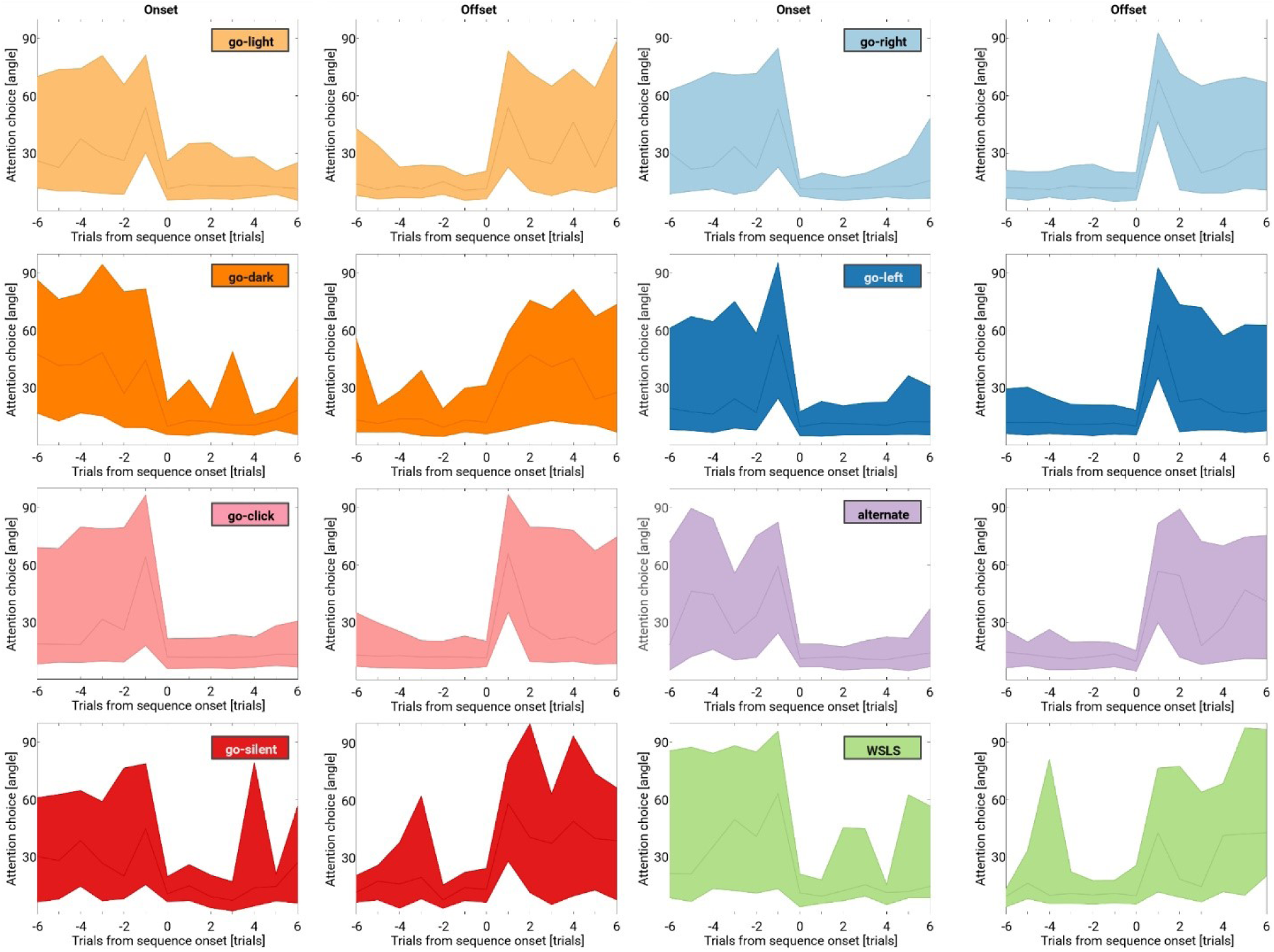
related to Figure 2. Head direction plots during choice formation for all strategies detected in naïve rats receiving random reward feedback. Similar plots as in Fig. S4 but this time the head direction plots are based on the average of multiple strategy sequences from 19 experimentally naïve rats performing eight sessions each with randomized reward feedback (i.e., without a task rule). This is interesting given that task feature-based learning in the absence of a clear behavioral advantage has also been reported in the context of human multidimensional learning.^4,5^ Statistics for head angle decrease/increase at strategy onset/offset are given below.

| Strategy | Number of sequences | Decrease in head angle at strategy onset | Increase in head angle at strategy offset |
| --- | --- | --- | --- |
| *alternate* | 75 | $3.7 \cdot 10^{-9}$ | $5.9 \cdot 10^{-9}$ |
| *go-click* | 432 | $1.7 \cdot 10^{-40}$ | $3.6 \cdot 10^{-47}$ |
| *go-dark* | 38 | $9.6 \cdot 10^{-3}$ | 0.03 |
| *go-left* | 353 | $1.6 \cdot 10^{-37}$ | $8.3 \cdot 10^{-42}$ |
| *go-light* | 102 | $8.9 \cdot 10^{-11}$ | $2.8 \cdot 10^{-11}$ |
| *go-right* | 135 | $1.1 \cdot 10^{-16}$ | $4.5 \cdot 10^{-17}$ |
| *go-silent* | 18 | $5 \cdot 10^{-3}$ | $8.6 \cdot 10^{-4}$ |
| *WSLS* | 38 | $9.9 \cdot 10^{-5}$ | $9.9 \cdot 10^{-5}$ |

**Figure S5.**
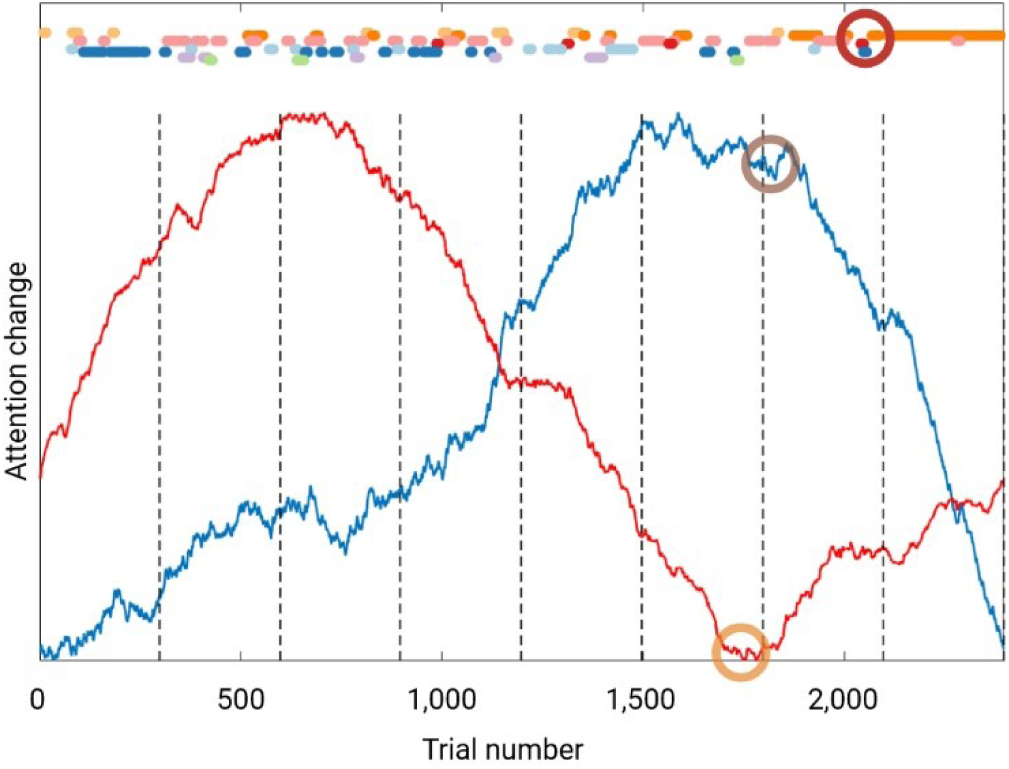
related to Figure 3. Attentional shifts across trials predict rule learning. Example rat learning the rule go-dark within eight sessions (dots represent trials, detected strategies are color-coded as in Figure 1). Dashed vertical lines mark the end of an experimental session. The empirical learning criterion (block length ≥ 30 trials) was reached after 2107 trials (red circle). Changes in attention during choice formation (blue curve) and reward feedback (red curve) are visualized using a CUSUM plot (cumulative sum of differences to the mean, Methods).^1^ Change points were detected using PARCS^6^ for attention-at-choice (grey circle: peak detected at trial 1833 corresponds to progressively smaller head angles between angle of view and correct side in the second after cue onset) and attention-at-reward (yellow circle: trough detected at trial 1778 corresponds to increasing angular sum caused by looking back-and-forth between food receptacle and side of chosen lever in the second after onset of reward feedback). Both change points were excellent predictors of rule learning at the group level (Figure 3I, Table S3).

**Table S1.** related to Figure 1. Strategy parameters for rats learning one rule in the multidimensional rule-learning task.

| Rule (n) | learn tresh [n] | y [t]** | perf cp [t]** | seq long [n]** | x<sub>1</sub> [t]** | x<sub>2</sub> [seq]* | x<sub>3</sub> [t]** | om [t]* |
| --- | --- | --- | --- | --- | --- | --- | --- | --- |
| Alternate (19) | 30 | 1,090 ± 116.8 | 957.3 ± 96.2 | 69 (52-97) | 459.5 ± 65.5 | 4.1 ± 0.6 | 1,150 ± 95 | 126 (25-256) |
| Go-click (17) | 58 | 462 ± 97.6 | 346.7 ± 77.11 | 116 (96-160) | 26.3 ± 14.4 | 6.5 ± 1.5 | n.a. | 6 (3-43.5) |
| Go-dark (24) | 30 | 1,336 ± 77.8 | 1,308 ± 66.3 | 191 (141.3-223.8) | 709.3 ± 86 | 4 ± 0.2 | 1,091 ± 58.6 | 18 (7-82.75) |
| Go-light (21) | 22 | 1,064 ± 105.4 | 1,115 ± 79.5 | 116 (79.5-207) | 259.8 ± 53.1 | 5 ± 0.5 | 998 ± 60.8 | 15 (3-108.5) |
| Go-right (11) | 42 | 74 ± 8.6 | 66.8 ± 8.3 | 181 (88-241) | 60.1 ± 5.8 | 0.6 ± 0.3 | n.a | 14 (2-80) |
| Go-silent (21) | 21 | 1,671 ± 136.4 | 1,628 ± 131 | 101 (69.5-138.5) | 1111 ± 127.7 | 3.7 ± 0.5 | 994.3 ± 95.3 | 15 (3-56) |
| WSLS (12) | 29 | 2,290 ± 307.2 | 2,008 ± 293 | 65 (52.25-90.5) | 776.5 ± 223.1 | 6.3 ± 1.4 | 3,214 ± 368.5 | 196 (48.75-336.8) |

The rule-specific empirical learning criterion is reached if the corresponding strategy sequence length exceeds *learn thresh*, *y* is the trial at which the empirical learning criterion is reached, *perf cp* is the trial at which the performance change point is reached, seq long is the longest correct strategy sequence (in trials), *x*_1_ is the trial when the correct strategy was followed for the first time, *x*_2_ is the number of sequences the correct strategy was followed prior to learning, *x*_3_ is a perseveration measure at the strategy level (measured in trials), *om* are the number of omissions.

**p<10-4, *p<2*10-3, one-way ANOVA or Kruskal-Wallis test (comparison of respective parameter between rules)

- *Learning & performance:* analyses were based on the empirical learning criterion and the performance change point (our two measures for learning) from 117/125 rats. n=8 rats (rule: win-stay-lose-shift/1, go-click/7) were excluded because either the empirical learning criterion was not reached (flat learning curve, n=2) or no significant performance change point (in most cases performance was high from beginning, n=6) could be detected. The longest correct sequence (i.e., the strategy that corresponds to the experimenter-defined rule) served as a measure of performance.
- Win-stay-lose-shift/WSLS: note that learning occurred in 11/12 rats (empirical learning criterion reached) but performance as measured by the parameter *longest correct sequence* was much lower as compared to other rules even though we trained rats for up to 20 days (we only stopped training if ≥50% of trials in a session were explained by detected *win-stay-lose-shift* sequences). Steady-state performance thus even stays lower with extensive training which may be at least partially explained by probabilistic feedback in this rule. Therefore, the detected performance change points are less meaningful than in other rules.
- The block length of *go-left* sequences corresponding to the empirical learning threshold for the rule go-left was 76 trials (this rule was only used in rats performing multiple switches).
- As a measure for strategy perseveration, we computed how many trials rats need to “sort out” the most salient strategy (defined as the 10-90%-rise time of the empirical cumulative distribution based on all detected sequences of that strategy). *Go-click* was chosen as the most salient strategy based on the finding that it was tested first across all rules (1 (1-2)). The only exception was the rule go-silent: in this case, we picked the *place* strategy the rat spontaneously used more often (*go-click* is less suitable here because it is the only strategy with consistent negative feedback and thus used less often). Due to the steep learning curve, values for the rules go-click and go-right are not meaningful and thus not reported.

**Table S2.** related to Figure 1. Linear regression model based on strategy measures.

Rats learning a single rule were included in this analysis. The linear regression model was fitted to predict the “learning trial” *y*(*R*) for rule *R* and based on 95/125 rats learning five different rules (alternate, go-dark, go-light, go-silent, win-stay-lose-shift). One rat (rule: win-stay-lose-shift) was excluded because it did not reach learning criterion, 29 rats (rule: one rat learning go-light, all rats learning the rules go-click & go-right) were excluded because the corresponding strategies were so salient for those rats that they almost directly selected them (i.e., in the absence of learning: first trial with correct strategy after 23 (5-61.5) trials). No outliers were removed. Results are provided for robust regression and ordinary least squares regression.

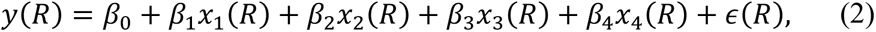

where,

- *x*_1_ is the trial when the correct strategy was followed for the first time.
- *x*_2_ is the number of sequences the correct strategy was followed prior to learning.
- *x*_3_ is a perseveration measure^2^ at the strategy level (Table S1).
- *x*_4_ is a categorical variable indicating the target rule.

The MATLAB routine *fitlm* transfers the categorical variable automatically into dummy indicator variables (1, when the rule is the target, experimenter-defined rule; 0, otherwise).

### Estimated Coefficients for robust regression

|  | Estimate | SE | tStat | pValue |
| --- | --- | --- | --- | --- |
| **(Intercept)** | -62.8 | 95.3 | -0.66 | 0.51 |
| **$x_1$** | 0.28 | 0.07 | 3.95 | $1.6 \cdot 10^{-4}$ |
| **$x_2$** | 77.52 | 11.07 | 7.00 | $5.0 \cdot 10^{-10}$ |
| **$x_3$** | 0.72 | 0.06 | 11.81 | $9.0 \cdot 10^{-20}$ |
| **$x_{4\_2}$** | 120.15 | 92.39 | 1.30 | 0.20 |
| **$x_{4\_4}$** | 415.37 | 106.8 | 3.89 | $2.0 \cdot 10^{-4}$ |
| **$x_{4\_7}$** | -108.2 | 94.26 | -1.15 | 0.25 |
| **$x_{4\_8}$** | -1267.8 | 163.05 | -7.78 | $1.4 \cdot 10^{-11}$ |

Number of observations: 95, Error degrees of freedom: 87, Root Mean Squared Error: 290, R^2^: 81.5%, Adjusted R^2^: 80.0%, F-statistic vs. constant model: 54.6, p-value = 3.7*10^-29^

### Estimated Coefficients for ordinary least squares regression

|  | Estimate | SE | tStat | pValue |
| --- | --- | --- | --- | --- |
| **(Intercept)** | 47.95 | 109.1 | 0.44 | 0.66 |
| **$x_1$** | 0.44 | 0.08 | 5.40 | $5.8 \cdot 10^{-7}$ |
| **$x_2$** | 102.73 | 12.68 | 8.10 | $3.1 \cdot 10^{-12}$ |
| **$x_3$** | 0.41 | 0.07 | 5.83 | $9.3 \cdot 10^{-8}$ |
| **$x_{4\_2}$** | 130.6 | 105.83 | 1.23 | 0.22 |
| **$x_{4\_4}$** | 353.79 | 122.34 | 2.89 | 0.0048 |
| **$x_{4\_7}$** | -45.53 | 107.97 | -0.42 | 0.67 |
| **$x_{4\_8}$** | -9.65 | 186.76 | -0.05 | 0.96 |

Number of observations: 95, Error degrees of freedom: 87, Root Mean Squared Error: 332, R^2^: 77.9%, Adjusted R^2^: 76.2%, F-statistic vs. constant model: 43.9, p-value = 6.4*10^-26^

**Table S3.** related to Figure 3. Linear regression model based on attention change points.

In a second regression model that used an independent data source based on head movements, we tested whether changes in strategy-specific attention markers during choice and reward feedback (as measured by the location of change points, Figure S5) also predict the “learning trial” *y*(*R*). n=29 rats performed 109 rules (excluding sessions with random reward feedback, either the sequence go-dark→ place→ alternate or go-dark→ place→ alternate→ go-silent).

### Attention-at-choice

we observed an orientation reaction after cue onset and tested whether sudden transitions (decrease across trials) in the minimum or the median angle of view with the respect to the side that corresponds to the correct strategy (in the second after cue onset) are detectable across trials. In 96/109 cases, we detected a significant change point for the minimum and in 90/109 cases for the median angle of view. We therefore used the minimum angle of view as a regressor in the model detailed below.

### Attention-at-reward

rats often looked back-and-forth between the pressed lever and the food receptacle during reward feedback. Likewise, we tested whether the angular sum in the second after the lever press significantly changes across trials (increase across trials). A change point was detected in 72/109 cases.

Correlation of the empirical learning criterion was high with both attention change points (R^2^ choice: 86.2%, R^2^ learning: 77.2%, Spearman correlation) and we included 65 rules from 29 rats for which both change points could be detected in our linear regression model. No regressor for rule type was included because it did not further improve results. No outliers were removed.

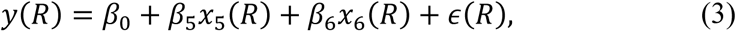

where,

- *x*_5_ = *CP*(*a*_AC_(*R*)) the change point trial of the angle-at-choice measurements.
- *x*_6_ = *CP*(*v*_AR_(*R*)) the change point trial of the attention-at-reward measurements.

### Estimated Coefficients for robust regression

|  | Estimate | SE | tStat | pValue |
| --- | --- | --- | --- | --- |
| (Intercept) | -11.93 | 43.65 | -0.27 | 0.79 |
| $x_5$ | 0.79 | 0.08 | 10.54 | $1.9 \cdot 10^{-15}$ |
| $x_6$ | 0.31 | 0.09 | 3.56 | $7.3 \cdot 10^{-4}$ |

Number of observations: 65, Error degrees of freedom: 62, Root Mean Squared Error: 148, R^2^: 91.7%, Adjusted R^2^: 91.5%, F-statistic vs. constant model: 343, p-value = 2.9*10^-34^

### Estimated Coefficients for ordinary least squares regression

|  | Estimate | SE | tStat | pValue |
| --- | --- | --- | --- | --- |
| (Intercept) | 27.35 | 57.08 | 0.48 | 0.63 |
| $x_5$ | 0.67 | 0.10 | 6.84 | $4.1 \cdot 10^{-9}$ |
| $x_6$ | 0.38 | 0.12 | 3.29 | $1.7 \cdot 10^{-3}$ |

Number of observations: 65, Error degrees of freedom: 62, Root Mean Squared Error: 193, R^2^: 85%, Adjusted R^2^: 84.5%, F-statistic vs. constant model: 175, p-value = 3*10^-26^

**Table S4.** Rat experimental groups used in this study.

- **n=17** rats were excluded (n=12 due to hardware/software problems or incomplete data, n=5 implanted rats excluded in the recovery period after surgery due to bad signal quality).
- The remaining **n=199** rats were distributed across experimental groups as follows:
- Conventional deterministic strategy set-shifting task **n=16** rats.
- Rats performing one rule in the multidimensional rule-learning task **n=125** rats (alternate n=19, go-click n=17, go-dark n=24, go-light n=21, go-right n=11, go-silent n=21, win-stay-lose-shift n=12). A separate cohort received random reward feedback (random **n=19**).
- Rats performing multiple rule switches in the multidimensional rule-learning task **n=33** rats (dark→ place→ alternate→ random: n=11; dark→ place→ alternate→ go-silent: n=22).
- Rats with silicon probes implanted into prelimbic prefrontal cortex **n=10** (n=5 probabilistic set-shift, n=5 multidimensional rule-learning task). For one rat performing the multidimensional rule-learning task, only four experimental days are available (the implant got damaged after the fourth session and we decided to sacrifice the animal to be able to determine electrode location). One implanted rat performing the probabilistic set-shift had difficulties to reach the performance threshold for rule switches (18 correct trials out of 20) and we therefore lowered the performance threshold to 16 correct trials out of 20 in later sessions. Since the electrophysiological focus of this study was on neural decoding of behavioral strategies (which are detected independent of task performance), no sessions from that animal were excluded for that reason.
- Video analyses were performed for **n=86** rats: all animals that received randomized reward feedback (n=19 experimentally naïve rats), for animals learning the rules alternate or go-silent only (n=19 rats each) and for 29 animals performing the multi-rule task (4 animals had incomplete video data).
- Note that the overall sum is 199 (instead of 203) rats because implanted rats performing multiple rule switches in the multidimensional rule-learning task were also used in behavioral analyses.

**Table S5.** Pseudorandomized list used for cue presentation in multidimensional rule-learning paradigm.

### Visual cue (0/1= left/right visual cue active)

1,0,1,0,1,0,1,0,1,1,1,0,0,1,1,0,0,1,0,0,1,1,0,0,0,1,1,0,1,0,1,0,0,1,1,0,0,1,1,0,0,1,0,1,0,1,1,1,0, 0,1,0,1,1,0,0,1,0,1,1,0,0,1,0,1,0,0,1,1,0,0,1,1,0,0,1,1,0,0,1,1,1,0,0,1,0,1,0,1,0,1,0,0,0,1,0,1, 0,1,1,0,1,0,0,1,1,0,0,1,1,1,0,0,0,1,1,0,0,0,1,1,0,1,1,0,1,0,1,1,0,0,1,0,0,1,1,0,1,1,0,1,0,0,1,0,1, 0,0,1,0,1,1,0,1,1,0,0,1,1,0,1,0,1,0,0,1,0,1,0,0,1,1,0,0,1,1,0,1,0,1,0,0,1,1,1,0,0,1,0,0,1,1,0,1, 0,1,1,0,1,0

### Auditory cue

0,1,1,0,0,1,1,1,0,1,0,0,1,0,1,0,1,1,0,1,0,1,1,0,1,0,1,1,1,0,0,0,1,1,0,0,1,1,0,1,0,0,1,1,0,0,1,0,0, 1,0,0,1,0,0,1,1,0,0,1,1,0,0,1,1,0,1,1,0,1,0,0,1,1,0,1,0,0,1,0,1,0,1,0,1,1,1,0,0,1,1,1,0,1,0,0,0, 1,1,0,0,1,1,0,0,1,1,0,1,0,1,0,1,0,0,1,0,1,0,1,0,0,1,0,0,1,1,0,1,0,1,1,1,0,0,1,1,0,1,1,0,0,1,1,0,0, 0,1,1,1,0,1,0,0,1,1,0,0,1,0,0,1,1,1,0,0,0,1,1,0,0,1,0,1,0,1,1,0,0,1,1,0,0,1,0,0,1,0,0,1,1,0,0,1, 1,0,1,0,0,1

For the multidimensional rule-learning task in rats and humans, we used a pseudorandomized list for cue presentation (instead of a truly randomized presentation) to avoid “accidental” reinforcement of non-sensory strategies in short blocks of trials during learning of a sensory rule. We took that measure because rats are prone to use *win-stay-lose-shift* or *place* strategies in operant chambers.^7^ The list always started at position one (and re-started there after 200 trials) and has the following properties: a cue is presented max. three times in a row on the same side, all sides (0 vs. 1/left vs. right with active cue) and cue pairs (visual-auditory: 0-1, 1-0, 0-1, 1-1) have the same frequency and no pair is directly repeated.

